# Neuroinvasion of SARS-CoV-2 in human and mouse brain

**DOI:** 10.1101/2020.06.25.169946

**Authors:** Eric Song, Ce Zhang, Benjamin Israelow, Alice Lu-Culligan, Alba Vieites Prado, Sophie Skriabine, Peiwen Lu, Orr-El Weizman, Feimei Liu, Yile Dai, Klara Szigeti-Buck, Yuki Yasumoto, Guilin Wang, Christopher Castaldi, Jaime Heltke, Evelyn Ng, John Wheeler, Mia Madel Alfajaro, Etienne Levavasseur, Benjamin Fontes, Neal G. Ravindra, David Van Dijk, Shrikant Mane, Murat Gunel, Aaron Ring, Syed A. Jaffar Kazmi, Kai Zhang, Craig B Wilen, Tamas L. Horvath, Isabelle Plu, Stephane Haik, Jean-Leon Thomas, Angeliki Louvi, Shelli F. Farhadian, Anita Huttner, Danielle Seilhean, Nicolas Renier, Kaya Bilguvar, Akiko Iwasaki

## Abstract

Although COVID-19 is considered to be primarily a respiratory disease, SARS-CoV-2 affects multiple organ systems including the central nervous system (CNS). Yet, there is no consensus whether the virus can infect the brain, or what the consequences of CNS infection are. Here, we used three independent approaches to probe the capacity of SARS-CoV-2 to infect the brain. First, using human brain organoids, we observed clear evidence of infection with accompanying metabolic changes in the infected and neighboring neurons. However, no evidence for the type I interferon responses was detected. We demonstrate that neuronal infection can be prevented either by blocking ACE2 with antibodies or by administering cerebrospinal fluid from a COVID-19 patient. Second, using mice overexpressing human ACE2, we demonstrate *in vivo* that SARS-CoV-2 neuroinvasion, but not respiratory infection, is associated with mortality. Finally, in brain autopsy from patients who died of COVID-19, we detect SARS-CoV-2 in the cortical neurons, and note pathologic features associated with infection with minimal immune cell infiltrates. These results provide evidence for the neuroinvasive capacity of SARS-CoV2, and an unexpected consequence of direct infection of neurons by SARS-CoV-2.

## Introduction

As of September 2020, SARS-CoV-2 has infected over 25 million people globally. While a majority of COVID-19 patients present with respiratory symptoms, neurological involvement, including impaired consciousness and headache, have been reported in patients (Mao et al., 2020). To date, human autopsy studies have identified viral RNA transcripts in brain tissues (Puelles et al., 2020; Solomon et al., 2020) and viral proteins in the endothelial cells within the olfactory bulb (Cantuti-Castelvetri et al., 2020) in a handful of people who succumbed to COVID-19. However, the potential of SARS-CoV-2 to infect neurons in the CNS is unknown. Understanding the full extent of viral invasion is crucial to treating patients, as we begin try to figure out the long-term consequences of COVID-19, many of which are predicted to have possible central nervous system involvement (De Felice et al., 2020; Heneka et al., 2020; Pereira, 2020; Zhang et al., 2020).

Because the central nervous system is not the primary organ affected by SARS-CoV-2, studying the neurological disease in COVID-19 patients systemically provides several challenges, including, having only a subset of the population of patients with neuroinvasion, lack of technology to sample CNS tissues directly, and being able to distinguish direct neuroinvasion vs. systemic viremia within the brain. Therefore, robust, reliable model systems are required to answer the questions underlying SARS-CoV-2 neuropathology. During the Zika virus (ZIKV) pandemic, several groups utilized human brain organoids to answer key questions regarding ZIKV neuroinvasion and its subsequent consequences (Garcez et al., 2016; Qian et al., 2016; Xu et al., 2019). Using a similarly well characterized human brain organoid model (Lancaster and Knoblich, 2014; Lancaster et al., 2013), here, we test the infection capacity of SARS-CoV-2 in central nervous system tissue. Utilizing single cell RNA-sequencing, we uncover the transcriptional changes caused by SARS-CoV-2 infection of neurons.

Beyond the neuroinvasive potential of SARS-CoV-2, the question remains whether or not ACE2 is the main route of entry of SARS-CoV-2 into neuronal cells and what strategies might block viral infection. ACE2 expression in the CNS, and in neurons in particular, is still unclear (Xia and Lazartigues, 2008). Moreover, in addition to ACE2, other cofactors such as TMPRSS2 (Hoffmann et al., 2020) and Neuropilin-1 (Cantuti-Castelvetri et al., 2020; Daly et al., 2020) seem to affect infection rates, but it is unclear if these are also required for neuronal infection. Thus, we use transcriptional profiling along with blocking antibody studies to demonstrate the requirement of ACE2 and SARS-CoV-2 spike protein to infect neurons.

To gain in vivo relevance of these findings, we examine the neuroinvasive potential SARS-CoV-2 using mouse models of SARS-CoV-2, and observe significant vascular remodeling in infected regions, independent of vascular infection, providing a valuable tool that can be used to dissect out the consequences of CNS infection. Finally, by examining post-mortem COVID-19 patient brain tissues, we provide evidence of neuroinvasion by SARS-CoV-2 and identify associations between infection and ischemic infarcts in localized brain regions.

## Results

### Modeling SARS-CoV-2 neuroinvasion and cellular death using human brain organoids

To dissect the mode and consequences of infection, we first established the neuroinvasive potential of SARS-CoV-2 in a human brain model system. We utilized human induced pluripotent stem cell (hiPSC) lines (Y1 and Y6), derived from healthy individuals (Figure S1), to generate forebrain-specific human neural progenitor cells (hNPCs). In culture, we observed replication of SARS-CoV-2 in two-week-old hNPCs, with peak viral titers as early as 12 h post infection (hpi) (Figure S2A and B). In addition, TUNEL staining indicated increased cell death (Figure S2A). Next, we generated hiPSC-derived brain organoids (Amin and Pasca, 2018; Pellegrini et al., 2020; Qian et al., 2016; Velasco et al., 2019) to model the SARS-CoV-2 infection of neuronal cells in 3D. Similar to a recent report (Ramani et al., 2020), we observed infection of neuronal cells in nine-week-old organoids as early as 24 hpi, with significantly increased number of SARS-CoV-2 positive cells at 96 hpi (Figure 1A-C, Figure S2C). Although the majority of the SARS-CoV-2 infected cells were localized within MAP2 positive cellular fields of mature neurons (Figure 1B.2, Figure S2C), we also observed infection of SOX2-positive neural stem cells with bipolar morphology and cells localized around the neural tube-like structures (Figure 1B.1). We observed an increase in SARS-CoV-2 positive cells 96 hpi compared to 24 hpi (Figure 1C, Figure S2C-E). At 96 hpi we observed a wide spread infection mostly limited to the regions with high cortical cell density in the organoid (Figure 1D). Using electron microscopy, we visualized viral particles within the organoid (Figure 1E, Figure S3), with discrete regions of high-density virus accumulation (Figure 1E.1) and other regions showing viral budding from endoplasmic reticulum-like structures (Figure 1E.2-4), suggesting SARS-CoV-2 is able to utilize the neuronal cell machinery to replicate. Organoid infection resulted in extensive neuronal cell death; strikingly, however, the majority of TUNEL-positive cells were SARS-CoV-2 negative (Figure 1F and G, Figure S2F), and only ∼15% of the cells infected with SARS-CoV-2 were TUNEL-positive (Figure 1H). Increased cell death was correlated with a higher density of SARS-CoV-2 positive cells (Figure 1I), which however, were not overtly dying (Figure 1G and H). Even within a single plane, we noticed high density SARS-CoV-2 area to have more TUNEL-positive cells (Figure 1J yellow box) compared to low-density SARS-CoV-2 regions (Figure 1J white box). Together, these data indicated that SARS-CoV-2 can infect cells of neural origin, and suggested that infected cells can promote death of nearby cells.

**Figure 1:**
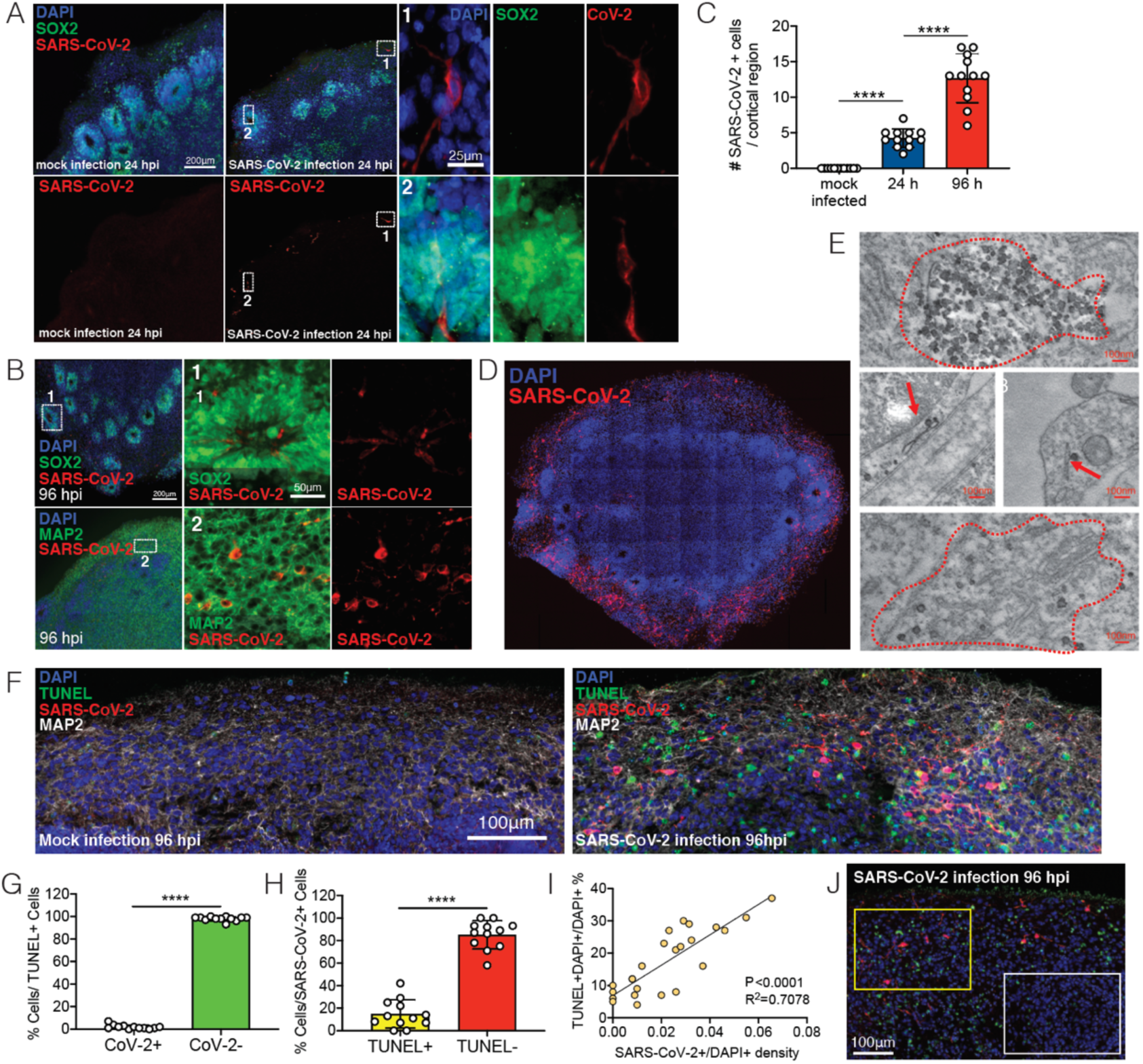
SARS-CoV-2 infects human brain organoids and induces cell death. Human brain organoids were infected with SARS-CoV-2 and collected at 24 hours post infection (hpi) or 96 hpi to analyze for different cellular markers. (**A**) Images of brain organoids looking at SARS-CoV-2 infection (in red) 24 hpi (See also Figure S2C for additional images). (**B**) Images of brain organoids looking at SARS-CoV-2 infection (in red) 96 hpi (See also Figure S2D for additional images). (**C**) Quantification of SARS-CoV-2 positive cells in a cortical region of organoids (**A-B**). (**D**) Tiled image of 96 hpi organoid. (**E**) Electron microscopy image of SARS-CoV-2 viral particles in brain organoids (See also Figure S3 for uncropped and additional images). (**F**) Organoids were stained with TUNEL to evaluate cell death at 96hpi. (**G**) Quantification of SARS-CoV-2 and TUNEL double positive (yellow) or SARS-CoV-2 negative, TUNEL positive (green) cells over total TUNEL positive cells. (**H**) Quantification of SARS-CoV-2 and TUNEL double positive (yellow) or SARS-CoV-2 positive, TUNEL negative (red) cells over total SARS-CoV-2 positive cells. (**I**) Correlation between the frequency of TUNEL positive cells and presence SARS-CoV-2 in different regions of the organoid. (**I**) Representative image of TUNEL and SARS-CoV-2 staining showing high-density SARS-CoV-2 region (yellow box) and low-density SARS-CoV-2 region (white box) in the same plane. All experiments were performed with unique organoid n of 4 per condition, from the same culturing batch, with images from n = 12 cortical regions with two IPSC lines, and student’s t-test was performed (****, P<0.0001).

### Single-cell level profiling of human brain organoid

We hypothesized that the cellular heterogeneity of brain organoids may be leading to certain cells being more susceptible to infection and others to death and that this system would provide an ideal platform to understand the cellular tropism of SARS-CoV-2 in the CNS and elucidate the consequences of SARS-CoV-2 neuroinvasion. To address these questions, we performed single-cell RNA sequencing to dissect the cellular states and transcriptional changes occurring after SARS-CoV-2 infection in both infected and non-infected cells within the organoid. We analyzed 60-day-old organoids that were either mock infected or infected with SARS-CoV-2 and collected at 2, 24, and 96 hpi (n = 2 for each time point); 96,205 cells were used to create 31 distinct clusters (Figure S4A). Annotations from previous studies (Cakir et al., 2019; Kanton et al., 2019; Velasco et al., 2019) allowed us to characterize the identity of each cluster (Figure S4A-C). Monocle trajectory further classified these clusters into four major cellular states; Neural progenitors/outer radial glia, intermediate progenitor/interneurons, neurons, and cortical neurons (Figure S4B-D). To confirm the cell-types we found in the single cell sequencing, we used additional staining with dorsal cortical markers CTIP2, PAX6, TBR1 and observed infection primarily overlapping with CTIP2 positive, PAX6 negative cells or CTIP2/TBR1 double positive, Pax6 negative cells, which are neuronal cells with a deep-layer (layers 5/6) fate.

Comparing global UMAP clusters in specimens collected before and after infection, we were able to identify cellular population changes during infection (Figure 2B, S4F and S5). With an added SARS-CoV-2 annotation, SARS-CoV-2 transcript reads were localized to a variety of cell clusters (Figure 2A and Figure S4G), demonstrating the widespread infectivity of SARS-CoV-2 in neurons, radial glia and neuronal progenitor cells. Several cell clusters—defined in Figure S4— showed large changes in their representation within the organoid, such as cluster 1 (from 5% to roughly 20% of the population), or cluster 7 (from ∼10% to nearly 0%) (Figure 2B). However, this was not uniformly seen with all infected clusters, as cells in cluster 11 showed high infection rates (Figure 2A), but with little change in population representation (Figure 2B, Figure S4H). This is consistent with findings from TUNEL staining (Figure 1E-G), which demonstrated minimal overlap between SARS-CoV-2 infected cells and those undergoing cell death.

**Figure 2:**
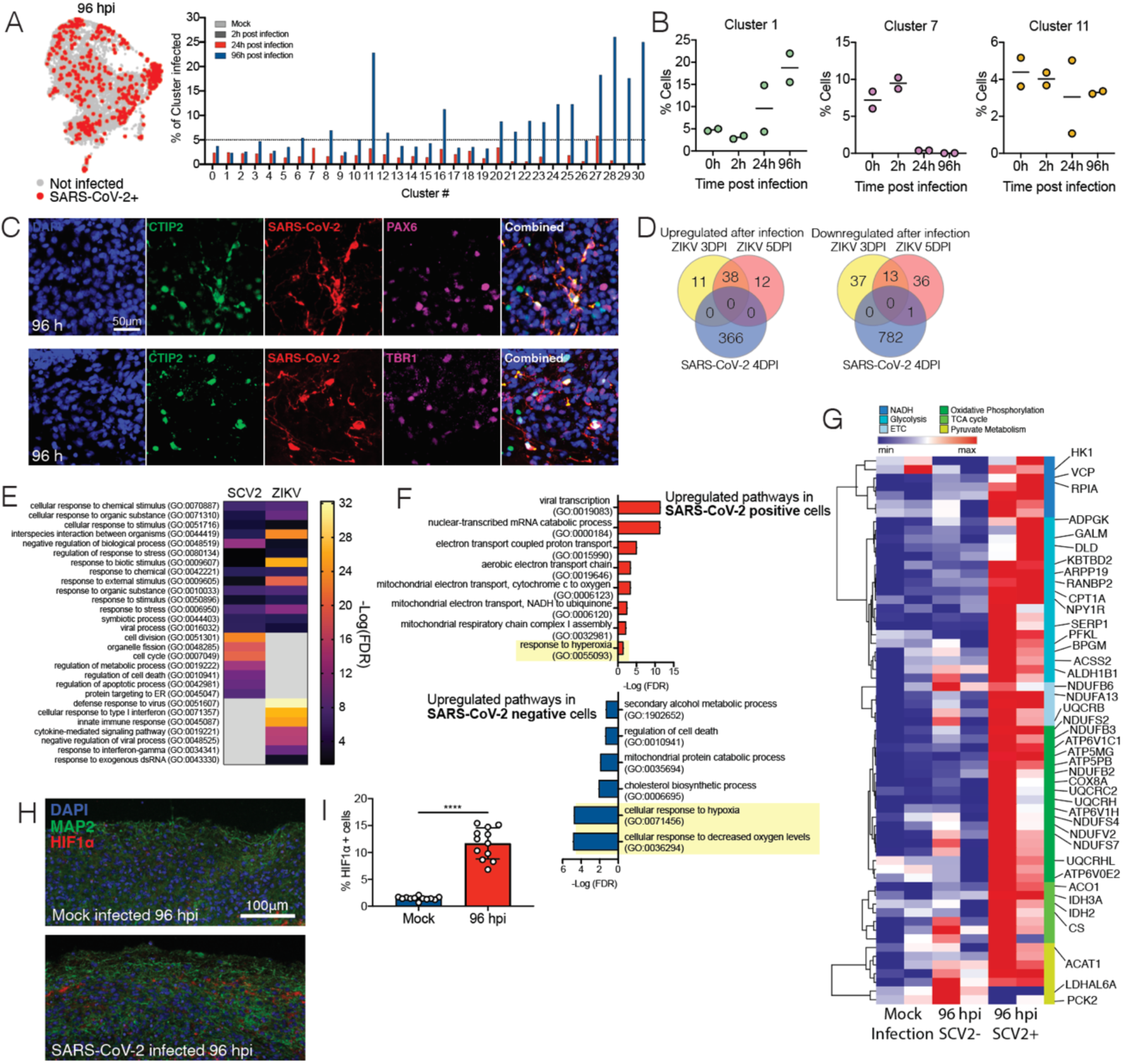
Neuronal cells undergo a unique metabolic response to SARS-CoV-2 infections. Brain organoids were infected with SARS-CoV-2 and sequenced with 10x single cell sequencing strategies. (**A**) Projection of cells with SARS-CoV-2 transcripts (in red) onto the UMAP. Percentage of infected cells in each cluster. (**B**) Change in population representation of example clusters after infection (refer See also Figure S5 for all clusters). (**C**) Validation of neuronal subtypes using CTIP2, PAX6 and TBR1 antibodies for confocal imaging. (**D**) Differentially expressed genes (DEGs) from brain organoids infected with ZIKV (Watnabe et al 2017 Cell Reports) were compared with DEGs from SARS-CoV-2 infected organoids. (**E**) Enriched gene ontology terms (geneontology.com) for upregulated genes from (**D**). (**F**) Enriched GO terms in SARS-CoV-2 infected cells (top) and SARS-CoV-2 negative bystander cells from 96 hpi organoids (bottom). (**G**) Heatmap of genes from metabolic pathways. (**H**) HIF1A staining of brain organoids that were mock infected versus 96hpi. (**I**) Quantification of HIF1A positive cells in SARS-CoV-2 infected organoids. Single cell RNA-seq was performed in duplicates with one IPSC line (Y6). HIF1A staining was performed with unique organoid n of 4 per condition, from the same culturing batch, with images from n = 12 cortical regions with two IPSC lines, and student’s t-test was performed (****, P<0.0001).

Next, we compared the changes in the transcriptome of SARS-CoV-2 infected organoids with another well-studied neurotropic virus—Zika virus (ZIKV). Differentially expressed genes (DEGs) after a brain organoid infection with ZIKV (Watanabe et al., 2017) showed almost no overlap with DEGs from SARS-CoV-2 infected organoids (Figure 2D). In both cases the transcriptome showed evidence of viral infection and invasion (Figure 2E). However, unique processes were enriched in each of the infections. SARS-CoV-2 infected brain organoid upregulated pathways related to cell division, organelle fission and metabolic processes, while ZIKV showed enrichment in type I interferon pathways (Figure 2E). SARS-Cov-2 induced a unique transcriptional state within neurons compared to ZIKV —consistent with the fact that SARS-CoV-2 induces a moderate interferon-stimulate gene (ISG) response in other tissues (Blanco-Melo et al., 2020), and previous reports of specific virus replication being controlled by alternative pathways by neurons (Daniels et al., 2019; Yordy et al., 2012).

To understand the impact of SARS-CoV-2 infection on the brain organoid at the cellular level, we characterized cells that were infected with SARS-CoV-2 versus neighboring uninfected cells without the presence of SARS-CoV-2 transcript, in a cluster comprising the highest number of infected cells at 96 hpi. SARS-CoV-2-positive cells showed enrichment of genes corresponding to viral transcription, along with enrichment for metabolic processes including electron transport coupled proton transport, cytochrome c to oxygen, and NADH to ubiquinone (Figure 2F top panel). Conversely, SARS-CoV-2-negative cells showed a mitochondrial catabolic state with the upregulation of alcohol metabolism, cholesterol synthesis, and regulation of cell death (Figure 2F bottom panel). These two cell populations displayed antagonizing pathway enrichment, with the infected cells responding to hyperoxia and the bystander cells responding to hypoxia (Figure 2F highlight in yellow). The hypermetabolic state is unique to the SARS-CoV-2 infected cells (Figure 2G) and highlights the ability of SARS-CoV-2 to hijack the host neuron machinery to replicate (Figure 1D). Finally, we confirmed that infection by SARS-CoV-2 induced a locally hypoxic environment in neuronal regions by staining for HIF1α (Figure 2H and 2I) in mock infected and SARS-CoV-2 infected organoids. Together these results indicate the potential of SARS-CoV-2 in manipulating host metabolic programming, that may create a resource-restricted environment for cells.

### Host ACE2 receptor is required for infection of neurons

One of the main arguments against SARS-CoV-2 neuroinvasion lies in the fact that the mRNA levels of ACE2 appear to be very low in the CNS (Li et al., 2020; Qi et al., 2020; Sungnak et al., 2020). Indeed, our single cell RNA-seq data set demonstrated low levels of ACE2; it was, however, detectable in many clusters (Figure S6A). In addition, we did not observe a correlation between the percentage of cells infected in each cluster to either ACE2, TMPRSS2 or Neuropilin-1 expression (Figure S6B). However, it remained possible that ACE2 protein may be expressed on the cell surface to promote viral entry. Consistent with this idea, we found widespread expression of ACE2 protein in both MAP2-positive neurons and cells in the neural tube-like structures of the organoids (Figure 3A), indicating that the mRNA level of ACE2 does not accurately reflect ACE2 protein expression. In addition, we used post-mortem human brain tissue to stain for neurons and ACE2 and found that neurons in the cortical grey matter co-localized with ACE2 staining (yellow arrow, Figure S6C), and found other cells in the vicinity that also stained for ACE2 (white arrow, Figure S6C). To test the requirement of ACE2 for SARS-CoV-2 infection, we incubated organoids with an anti-ACE2 blocking monoclonal antibody prior to infection with SARS-CoV-2. We detected significant inhibition of SARS-CoV-2 infection upon pre-treatment with ACE2 antibody compared with isotype control, indicating the requirement of ACE2 for infection of brain organoids (Figure 3B and 3F, Figure S6C).

**Figure 3:**
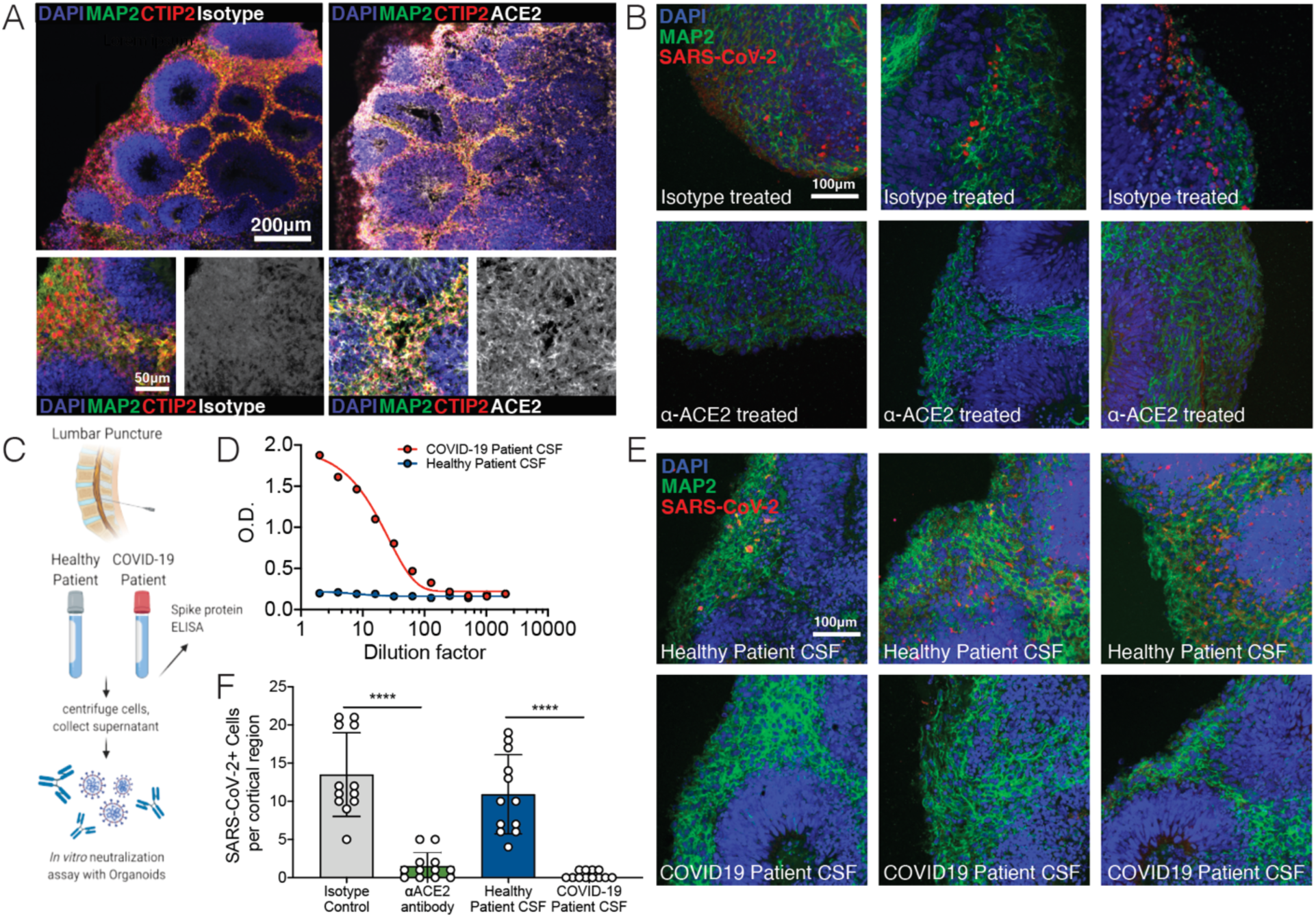
SARS-CoV-2 neural infection depends on ACE2 and can be neutralized by anti-spike antibodies found in CSF of COVID-19 patients. (**A**) Immunofluorescence staining of ACE2 in brain organoids. (**B**) Immunofluroescence staining of organoids pre-incubated with isotype antibodies (top row) or anti-ACE2 antibodies (bottom row) and infected with SARS-CoV-2. (**C**) Schematic showing collection of clinical lumbar puncture from patients with and without COVID-19 for assays shown in (**D-F**). (**E**) Immunofluroescence staining of organoids infected with SARS-CoV-2 preincubated with CSF from health patients (top row) or CSF from COVID-19 patients (bottom row). (**F**) Quantification of figures from (**C** and **E**). All experiments were performed with unique organoid n of 4 per condition, from the same culturing batch, with images from n = 12 cortical regions with two IPSC lines, and student’s t-test was performed (****, P<0.0001).

Next, we investigated whether there are humoral antibody responses against SARS-CoV-2 in the CNS of infected patients, and whether antibodies present in the CNS can prevent infection of neurons. We analyzed cerebrospinal fluid (CSF) from a patient hospitalized with COVID-19 and acute encephalopathy (Figure 3C) and from a healthy control volunteer by performing ELISA against the spike protein of SARS-CoV-2 (Figure 3D). We detected IgG antibodies specific to the spike protein in the patient CSF even at 100x dilutions (Figure 3D). Using this patient’s CSF, we performed a neutralization assay against SARS-CoV-2 and validated its efficacy in preventing brain organoid infection (Figure 3E). CSF-containing antiviral antibodies blocked SARS-CoV-2 infection in organoids (Figure 3E and F, Figure S6D), highlighting the potential of SARS-CoV-2 neuroinvasion and subsequent immune response in the CNS.

### Mouse models of COVID-19 confirm the neuroinvasive potential of SARS-CoV-2

In order to examine the consequences of SARS-CoV-2 infection in a more physiologically complete system, we examined the neuroinvasive potential of SARS-CoV-2 *in vivo* by using transgenic mice expressing human ACE2 under the K18 promoter (K18-hACE2) (McCray et al., 2007). Similar to previous reports of SARS-CoV showing neurotropism (McCray et al., 2007; Netland et al., 2008), we observed increasing viral titers in the brain of mice after intranasal administration of SARS-CoV-2 (Figure 4A and B). We next analyzed the distribution of the virus in the whole brain by immunolabeling against the nucleocapsid protein, clearing and light sheet microscopy imaging using iDISCO+ (Renier et al., 2014) (Figure 4, Supplementary Movie 1). At 7 days after infection, the virus was widely present in neural cells throughout the forebrain (Figure 4C). The cortex was unevenly infected, as the infected cells were visible in columnar patches and in sensory regions, while the layer 4 was mostly devoid of infection (Figure 4C and D). We mapped the density of infected cells using ClearMap (Renier et al., 2016) and confirmed that most brain regions contained high density of infected cells, to the notable exception of the cerebellum (Figure 4E). Other regions also contained relatively low density of infected cells, for instance the dentate gyrus, the globus pallidus and cortical layer 4. Of note, the whole brain analysis of the virus distribution didn’t detect the presence of the virus in the vascular endothelium. To explore the possibility that the viral expression by neural cells could indirectly affect the organization of the vascular network, we double labeled brains for the nucleocapsid protein and the vascular endothelium with CD31 and Podocalyxin, and used again ClearMap to reconstruct the whole brain vascular network 7 days after infection (Figure 4F) (Kirst *et al*. Cell 2020). We focused on the organization of the cortical vasculature. In infected brains, we focused on hot spots of infections represented as the number of detected infected cells and measured the density and orientation of all blood vessels. In many instances, the expression of the virus coincided with a disruption in the normal vascular topology expected in the cortex, with an important loss of the normal enrichment in radially oriented vessels in upper layers. This analysis suggests that an important vascular remodeling accompanies the expression of the virus by neural cells, presumably reorienting the normal blood flow towards metabolically active hot spots.

**Fig. 4:**
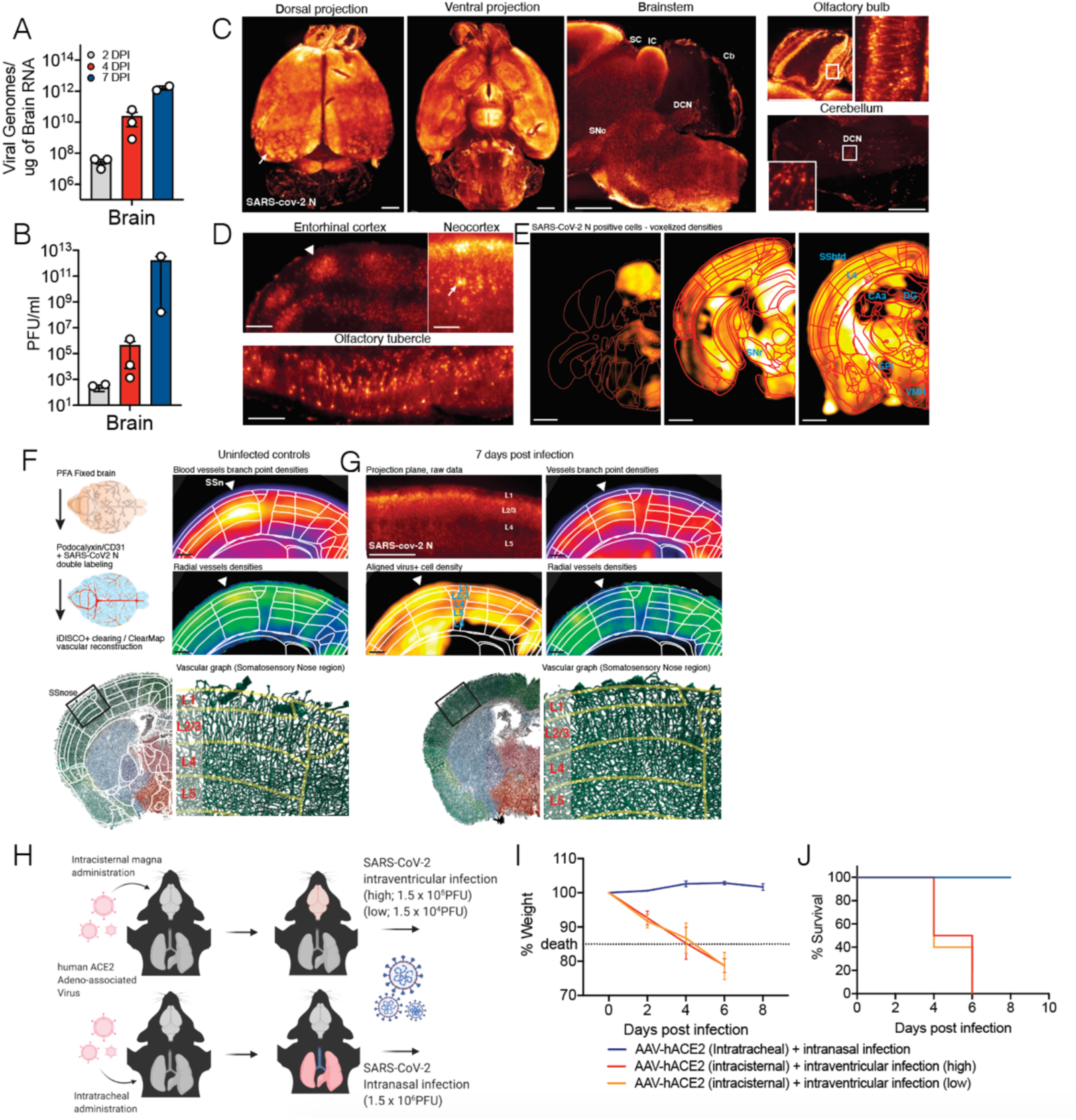
SARS-CoV-2 replicates efficiently in the brain of mice and can cause central nervous system specific lethality. (**A-C**) Mice expressing human ACE2 under the K18 promoter (K18-hACE2) were infected with SARS-CoV-2 intranasally, and brains of the mice were collected at days 2, 4 and 7 post infection for (**A**) qPCR or (**B**) plaque assay. (**C-E**) iDISCO+ whole brain immunolabeling against the nucleocapsid protein of SARS-CoV2 7 days after an intranasal infection, shown as 300µm projection planes. (**C**) Dorsal, ventral and sagittal projections showing widespread distribution of the virus in the forebrain with patches of high viral density in the cortex (arrow). The virus is not detected in the cerebellum, except for the pial meninges and DCNs. (**D**) 300µm deep projection planes in the cortex showing cortical patches of viral expression (arrowhead), a reduced infection of the cells in the layer 4, and expression in pyramidal neurons (arrow). (**E**) ClearMap analysis of the infected cells distribution (n=3), registered to the Allen Brain Atlas, showing the wide distribution of the virus across brain regions, with a few regions with lower densities, among which the DG, GPi, CA3, cortical layer 4 and VMH. (**F-G**) Analysis of the vascular network using ClearMap and iDISCO+ 7 days after intranasal infection by mapping of the vascular network with a co-labeling of the N protein. Planes at the level of the Nose somatosensory cortex are shown. (**F**) Control uninfected brains. Branch point densities (top panel) peak in controls at layer 4. The density of radially oriented vessels (middle panel) peaks in layers 1,2 and 3 while decreasing in the layers 4, 5 and 6. (**G**) 7 days post-infection brain. Expression of the N viral protein by neural cells are shown at the level of the nose somatosensory cortex (300µm projection plane and mapped densities). While branch point densities of vessels still show a peak in layer 4, the normal radial organization of the vessels is not measured in the Nose region (arrowhead). Representative render of the vascular graph are showing a decrease in the vessels orientations seen in the control in layers 2 and 3.(**H**) Schematic of experiment for (**I-J**), adeno-associated virus coding for human ACE2 (AAV-hACE2) were injected in to the cisterna magna or intratracheally to induce brain specific or lung specific expression of hACE2. Brain hACE2 expressing mice were infected with SARS-CoV-2 intraventricularly, and lung hACE2 expressing mice were infected with SARS-CoV-2 intranasally. (**I**) Weight loss curve and (**J**) survival curve of mice infected with SARS-CoV-2 in the lung (blue) and the brain (red and orange) (blue, n=10; red, n=4; orange, n=4). Scale bars are 1mm (A,C), 200µm (B) and 500µm (D,E) Abbreviations: CB: Cerebellum, DCN: Deep Cerebellar Nuclei, GPi: Globus Pallidus internal segment, IC: Inferior Colliculus, SC: Superior Colliculus, SN: Substantia Nigra (reticulata or compacta), SS: Somatosensory Cortex, Nose of Barrel FielD, VMH: Ventro Medial Hypothalamus

To dissect the consequences of SARS-CoV-2 infection in the CNS versus the respiratory system, we directed expression of human ACE2 using an adeno-associated virus vector (AAV-hACE2) (Israelow et al., 2020) to the lungs via intratracheal (IT) delivery or to the brain via intracisternal (IC) delivery (Figure 4H). Then, the mice were infected with SARS-CoV-2 respectively, either intranasally or intraventricularly. Intranasally infected mice showed signs of lung pathology (Israelow et al., 2020) but no weight loss or death (Figure 4I and J). However, intraventricular administration of SARS-CoV-2 resulted in significant weight loss and death, even at the challenge virus dose 100-fold lower than that used for intranasal infection (Figure 4I and J). Altogether, this highlights the neuro-replicative potential and lethal consequences of SARS-CoV-2 CNS infection in mice.

### Evidence of SARS-CoV-2 neuroinvasion in COVID-19 patient brain autopsies

Finally, to determine whether there is evidence of SARS-CoV-2 CNS infection in COVID-19 patient, we examined paraffin embedded formalin fixed sections of three patients who died after suffering from severe COVID-19-related complications. All patients had been admitted to the ICU, they were sedated and ventilated due to respiratory failure (days 3, 10 and 18; patient 3, 1 and 2 respectively), and their difficulty to be weaned from mechanical ventilation indicated the severity and highly pathogenic nature of the disease course (Supplementary Table 1). We sampled several regions of the brain and stained for both, Hematoxylin and Eosin (H&E) and for SARS-CoV-2 spike protein with a validated protocol and antibody we have used previously (Hosier et al., 2020). While in control slides there was no positive immunohistochemical staining for the SARS-CoV-2 spike protein, samples from COVID-19 patients showed distinct, and specific spike protein staining, albeit to different degrees (Figure 5, Figure S7).

**Figure 5:**
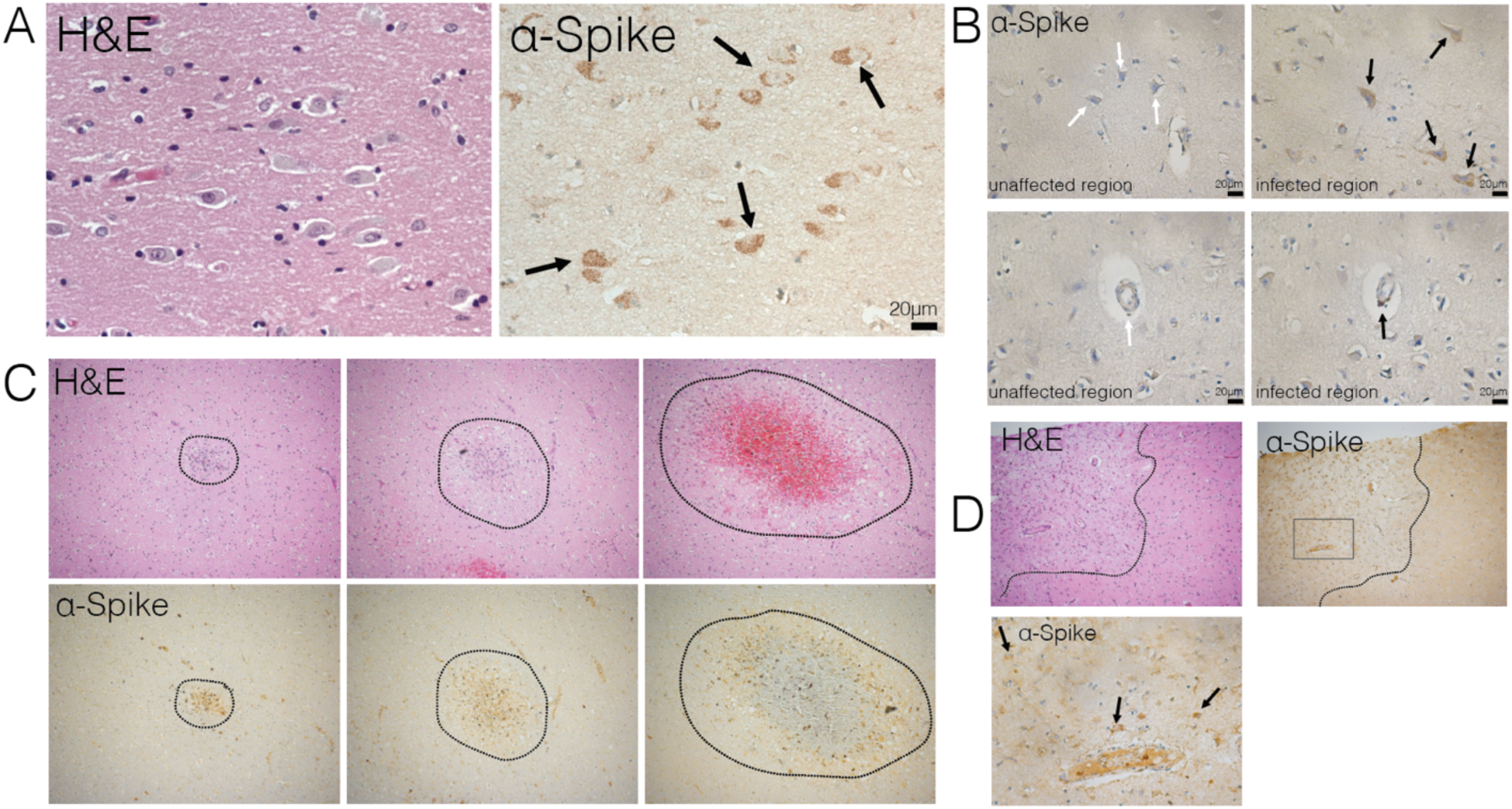
Evidence of neuroinvasion in post-mortem COVID-19 patient brains. FFPE sections of brain tissue from COVID-19 patients were stained using H&E and anti-SARS-CoV-2-spike antibody. (**A**) Image of cortical neurons positive for SARS-CoV-2 (black arrows). (**B**) Images of unaffected regions (left) and infected regions (right) demonstrating infection of neurons (top row) and microvasculature (bottom row). (**C**) Ischemic infarcts found at different stages stained with H&E (top row) and SARS-CoV-2-spike antibody (bottom row). (**D**) Ischemic region outlined with dotted line with positive staining focused around ischemic infarct. Bottom image shows zoomed in image indicated by dotted box in top image, and black arrows indicate infected neurons in the region.

Neuropathological evaluations identified staining with anti-spike antibodies in cortical neurons (Figure 5A, B upper panel, D black arrows) along with signal in endothelial cells (Figure 5B lower panel) in patient 1 medium spiny neurons in the basal ganglia, however, did not show staining, similar to substantia nigra neurons in the midbrain which were also negative for the SARS-CoV-2 spike protein (Figure S8). The cellular staining pattern showed diffuse cytoplasmic and perinuclear staining, along with concentrated regions within the cells—consistent with our findings using electron microscopy in the organoids, which demonstrated localized hotspots within the infected cell. Patient 3, who failed to regain consciousness after cessation of sedation, was diagnosed with severe global encephalopathy and was noted with multiple diffuse micro bleeds as demonstrated by diagnostic imaging (MRI). Upon histologic examination, we found multiple microscopic ischemic infarcts in the subcortical white matter, ranging from acute to subacute, and with focal hemorrhagic conversion (Figure 5C, Figure S9A). These varying stages of infarction indicated a temporal sequence of continued ischemic events. Most infarcts showed signs of tissue damage and localized cell death, and positive viral staining was present predominately around the edges of the infarct and to a lesser degree within the center. At the hyperacute stage of infarction, viral proteins were present in endothelium. These infarcts are present predominately in subcortical white matter, not within the cortex. In other samples, we found suspected viral staining in locally ischemic regions (Figure 5D, Figure S9B and C). Remarkably, all the regions of positive viral staining showed no lymphocyte or leukocyte infiltration. This is in contrast to other neurotropic viruses—Zika virus, rabies virus, herpes virus— in which the infection is typically accompanied by large number of immune cell infiltrates, including T cells. These findings suggest that, although SARS-CoV-2 has neurotropic properties and can infect neurons in patients, it did not invoke an immune response typical of other neurotropic virus in this particular case.

## Discussion

We examined the potential for SARS-CoV-2 to infect neural tissues of both mice and human origin and demonstrate potential consequences of its neuroinvasion. Our results suggest that neurologic symptoms associated with COVID-19 may be related to consequences of direct viral invasion of the CNS. Specifically, our work experimentally demonstrates that the brain is a site for high replicative potential for SARS-CoV-2. We further show that SARS-CoV-2 causes significant neuronal death in human brain organoids. Using electron microscopy, we identified viral particles budding from the endoplasmic reticulum, indicating the virus’s ability to use the neuron cell machinery to replicate. Similar to neuronal loss observed in patient autopsies (Solomon et al., 2020), we noticed large numbers of cells dying in the organoid, however, this neuronal death did not colocalize directly with virus infection. Single-cell RNA sequencing of the infected organoids showed metabolic changes in neurons without interferon or ISG signatures, indicating the neuroinvasive consequence of SARS-CoV-2 is unique compared to other neurotropic viruses such as ZIKV. Closer examination showed diverging metabolic changes in infected versus neighboring cells, suggesting that the infected cells can cause local changes to their microenvironment affecting survival of nearby cells. It is possible that viral infection induces locally hypoxic regions which aids in lowering the threshold for tissue damage in the context of an already oxygen deprived state.

While ACE2 expression levels in the human brain are still being investigated, we showed that ACE2 is expressed at the protein level and is functionally required for SARS-CoV-2 infection in human brain organoids. Further, we detected robust antiviral antibody presence in the CSF of a COVID-19 patient, who presented with acute neurologic symptoms. This finding suggests that, at least in some patients with COVID-19 and neurological symptoms, there is robust antibody response against the virus within the CSF. Although we did not find induction of ISGs in brain organoids, or leukocyte infiltration in the post-mortem COVID-19 brain tissues, the neuroinflammatory consequences of SARS-CoV-2 cannot be excluded given the amount of cell death we observed. In the in vivo setting of the CNS with vasculature and immune cells, neuronal death could have cascading downstream effects in causing and amplifying CNS inflammation.

Although our rodent model does not utilize the endogenous ACE2 expression, it has been previously reported that even mouse adapted SARS-CoV is still neurotropic in wild type mice and SARS-CoV-2 is neurotropic in mice with hACE2 expression from the endogenous locus (Roberts et al., 2007; Sun et al., 2020). Using mouse models, we demonstrate for the first time that SARS-CoV-2 neuroinvasion in mice can have significant remodeling of brain vasculature, providing a link between the hypoxia and what we see in both the human organoid and the patient brains.

Similar to previous reports of acute hypoxic ischemic damage without microthrombi in post-mortem brain of COVID-19 patients (Solomon et al., 2020), we also found presence of ischemic damage and micro infarcts in our post-mortem brain samples of COVID-19 patients. In our study, we observed evidence of SARS-CoV-2 infection within the regions of micro ischemic infarcts suggesting the possibility of neuroinvasion associated ischemia and vascular anomalies, consistent with what we observe in mice. However, a limitation of our study is that autopsy samples from only a small number of patients were examined. Future studies are needed to examine whether there are other cases of neuroinvasion in the CNS, and the predisposition for such infection. Although we are unable to determine the exact relationship between neuroinvasion and ischemic infarcts, we pose a possible hypothesis from our findings in the patients, mice, and infections of human brain organoids; that SARS-CoV-2 neuroinvasion may cause locally hypoxic regions and disturbance of vasculature, and the disruption of brain vasculature can make vulnerable ischemic infarcts and regions more susceptible to viral invasion (Figure S10). Our findings expand the utility of human brain organoids, beyond modeling fetal brains, and highlight the importance of using a variety of approaches to best model physiology of the human brain.

In future studies, identifying the route of SARS-CoV-2 invasion into the brain in addition to determining the sequence of infection in different cell types in the central nervous system will help validate the temporal relationship between SARS-CoV-2 and ischemic infarcts in patients. It may be through the nasal cavity to CNS connection through the cribriform plate, olfactory epithelium and nerve, or viremia, but regardless, the brain should be considered a high-risk organ system upon respiratory exposure (Baig and Sanders, 2020; Coolen et al., 2020).

Altogether, our study provides clear demonstration that neurons can become a target of SARS-CoV-2 infection, with devastating consequences of localized ischemia in the brain and cell death, highlighting SARS-CoV-2 neurotropism and guiding rational approaches to treatment of patients with neuronal disorders.

All procedures were performed in a BSL-3/ABSL-3 facility (for SARS-CoV-2-related work) with approval from the Yale Environmental Health and Safety committee (#20-19 and #18-16).

## Mice

Six to twelve-week-old mixed sex C57Bl/6 (B6J) purchased from Jackson laboratories, and B6.Cg-Tg(K18-ACE2)2Prlmn/J(K18-hACE2) mice (gift from Jackson Laboratories) were subsequently bred and housed at Yale University. All procedures used in this study (sex-matched, age-matched) complied with federal guidelines and the institutional policies of the Yale School of Medicine Animal Care and Use Committee.

## AAV infection (Intratracheal and Intracisternal magna injection)

Adeno-associated virus 9 encoding hACE2 were purchased from Vector biolabs (AAV-CMV-hACE2).

### Intratracheal injection

Animals were anaesthetized using a mixture of ketamine (50 mg kg^−1^) and xylazine (5 mg kg^−1^), injected intraperitoneally. The rostral neck was shaved and disinfected. A 5mm incision was made and the salivary glands were retracted, and trachea was visualized. Using a 500μL insulin syringe a 50μL bolus injection of 10^11^GC of AAV-CMV-hACE2 was injected into the trachea. The incision was closed with VetBond skin glue. Following intramuscular administration of analgesic (Meloxicam and buprenorphine, 1 mg kg^−1^), animals were placed in a heated cage until full recovery.

### Intracisternal magna injection

Mice were anesthetized using ketamine and xylazine, and the dorsal neck was shaved and sterilized. A 2 cm incision was made at the base of the skull, and the dorsal neck muscles were separated using forceps. After visualization of the cisterna magna, a Hamilton syringe with a 15 degree 33 gauge needle was used to puncture the dura. 3μL of AAV_9_ (3.10^12^ viral particles/mouse) or mRNA (4-5 μg) was administered per mouse at a rate of 1μL min^-1^. Upon completion of the injection, needle was left in to prevent backflow for an additional 3 minutes. The skin was stapled, disinfected and same post-operative procedures as intratracheal injections were performed.

## Generation of SARS-CoV-2 virus

To generate SARS-CoV-2 viral stocks, Huh7.5 cells were inoculated with SARS-CoV-2 isolate USA-WA1/2020 (BEI Resources #NR-52281) to generate a P1 stock. To generate a working stock, VeroE6 cells were infected at a MOI 0.01 for four days. Supernatant was clarified by centrifugation (450g x 5min) and filtered through a 0.45 micron filter. To concentrate virus, one volume of cold (4 °C) 4x PEG-it Virus Precipitation Solution (40% (w/v) PEG-8000 and 1.2M NaCl) was added to three volumes of virus-containing supernatant. The solution was mixed by inverting the tubes several times and then incubated at 4 °C overnight. The precipitated virus was harvested by centrifugation at 1,500 x g for 60 minutes at 4 °C. The pelleted virus was then resuspended in PBS then aliquoted for storage at -80°C. Virus titer was determined by plaque assay using Vero E6 cells.

## SARS-CoV-2 infection of organoids

Brain organoids in low adhesion plates were infected with SARS-CoV-2 at a MOI of 1.

## SARS-CoV-2 infection (intranasal)

Mice were anesthetized using 30% v/v Isoflurane diluted in propylene glycol. Using a pipette, 50μL of SARS-CoV-2 (3×10^7^ PFU/ml) was delivered intranasally.

## SARS-CoV-2 infection (intraventricular)

Animals were anaesthetized using a mixture of ketamine (50 mg kg^−1^) and xylazine (5 mg kg^−1^), injected intraperitoneally. After sterilization of the scalp with alcohol and betadine, a midline scalp incision was made to expose the coronal and sagittal sutures, and a burr holes were drilled 1 mm lateral to the sagittal suture and 0.5 mm posterior to the bregma. A 10 μl Hamilton syringe loaded with virus, and was inserted into the burr hole at a depth of 2 mm from the surface of the brain and left to equilibrate for 1 min before infusion. Once the infusion was finished, the syringe was left in place for another minute before removal of the syringe. Bone wax was used to fill the burr hole and skin was stapled and cleaned. Following intramuscular administration of analgesic (Meloxicam and buprenorphine, 1 mg kg^−1^), animals were placed in a heated cage until full recovery. For high condition, 5μL of SARS-CoV-2 (3×10^7^ PFU/ml) and for low condition 5μL of SARS-CoV-2 (3×10^6^ PFU/ml) was used.

## Samples staining and iDISCO+ clearing

Whole brain vasculature staining was performed following the iDISCO+ protocol previously described (Renier et al., 2016) with minimal modifications. All the steps of the protocol were done at room temperature with gentle shaking unless otherwise specified. All the buffers were supplemented with 0,01% Sodium Azide (Sigma-Aldrich, Germany) to prevent bacterial and fungi growth. Brains were dehydrated in an increasing series of methanol (Sigma-Aldrich, France) dilutions in water (washes of 1 hour in methanol 20%, 40%, 60%, 80% and 100%). An additional wash of 2 hours in methanol 100% was done to remove residual water. Once dehydrated, samples were incubated overnight in a solution containing a 66% dichloromethane (Sigma-Aldrich, Germany) in methanol, and then washed twice in methanol 100% (4 hours each wash). Samples were then bleached overnight at 4°C in methanol containing a 5% of hydrogen peroxide (Sigma-Aldrich). Rehydration was done by incubating the samples in methanol 60%, 40% and 20% (1 hour each wash). After methanol pretreatment, samples were washed in PBS twice 15 minutes and 1 hour in PBS containing a 0,2% of Triton X-100 (Sigma-Aldrich) and further permeabilized by a 24 hours incubation at 37°C in *Permeabilization Solution*, composed by 20% dimethyl sulfoxide (Sigma-Aldrich), 2,3% Glycine (Sigma-Aldrich, USA) in PBS-T. In order to start the immunostaining, samples were first blocked with 0,2% gelatin (Sigma-Aldrich) in PBS-T for 24 hours at 37°C, the same blocking buffer was used to prepare antibody solutions. A combination of primary antibodies targeting different components of the vessel’s walls were used to achieve continuous immunostaining. Antibodies to Podocalyxin and CD31 were combined with antibodies against the nucleocapsid (N) from GeneTex (Antibodies’ references and concentrations are provided in Sup. Table). Primary antibodies were incubated for 10 days at 37°C with gentle shaking, then washed in PBS-T (twice 1 hour and then overnight), and finally newly incubated for 10 days with secondary antibodies. Secondary antibodies conjugated to Alexa 647 were used to detect Podocalyxin and CD31, while the Nucleocapsid protein was stained with a secondary antibody conjugated to Alexa 555. After immunostaining, the samples were washed in PBS-T (twice 1 hour and then overnight), dehydrated in a methanol/water increasing concentration series (20%, 40%, 60%, 80%, 100% one hour each and then methanol 100% overnight), followed by a wash in 66% dichloromethane – 33% methanol for 3 hours. Methanol was washed out with two final washes in dichloromethane 100% (15 min each) and finally the samples were cleared and stored in dibenzyl ether (Sigma-Aldrich) until light sheet imaging.

## Light sheet imaging

We imaged with a 4X 0.35NA objective cropped elongated field of view (600 × 2200µm) covering the narrow waist of the light sheet at 1.63µm/pixel of lateral resolution at 1.6µm spacing. A reference channel for the registration to the annotated atlas using the sample autofluorescence was acquired at 5µm/pixel. The acquisitions were done on a LaVision Ultramicroscope II equipped with infinity-corrected objectives. The microscope was installed on an active vibration filtration device, itself put on a marble compressed-air table. Imaging was done with the following filters: 595/40 for Alexa Fluor-555, and -680/30 for Alexa Fluor-647. The microscope was equipped with the following laser lines: OBIS-561nm 100mW, OBIS-639nm 70mW, and used the 2nd generation LaVision beam combiner. The images were acquired with an Andor CMOS sNEO camera. Main acquisitions were done with the LVMI-Fluor 4X/O.3 WD6 LaVision Biotec objective. The brain was positioned in sagittal orientation, cortex side facing the light sheet. A field of view of 400 × 1300 pixels was cropped at the center of the camera sensor. The light sheet numerical aperture was set to the NA (0.1). Beam width was set to the maximum. Only the center-left light sheet was used. Laser powers were set to 100% (639nm) or 10% (561nm). Tile overlaps were set to 10%. The acquisition routine was set to 1) Z-drive -> Save ome.tif stack 2) Filter change -> Z-drive -> Save ome.tif stack 3) Change X position -> repeat 1,2 12 times 4) Change y position -> repeat 1,2,3 6 times. At the end of the acquisition, the objective is changed to a MI PLAN 1.1X/0.1 for the reference scan at 488nm excitation (tissue autofluorescence). The field of view is cropped to the size of the brain, and the z-steps are set to 6µm, and light sheet numerical aperture to 0.03 NA.

## Viral RNA analysis

At indicated time points mice were euthanized in 100% Isoflurane. Brain tissue was placed in a bead homogenizer tube with 1ml of PBS+2%FBS. After homogenization 250ul of this mixture was placed in 750ul Trizol LS (Invitrogen), and RNA was extracted with RNeasy mini kit (Qiagen) per manufacturer protocol. To quantify SARS-CoV-2 RNA levels, RT-qPCR was performed using Luna Universal Probe Onestep RT-qPCR kits (New England Biolabs) with 1 ug of RNA, with the US CDC real-time RT-PCR primer/probe sets for 2019-nCoV_N1.

## Viral titer

Brain homogenates were centrifuged at 3900g for 10 minutes and supernatant was taken for plaque assays. Supernatant at limiting dilutions were incubated on Vero E6 cells in MEM supplemented NaHCO_3_, 4% FBS 0.6% Avicel RC-581. Plaques were resolved at 48hrs post infection by fixing in 10% formaldehyde for 1 hour followed by staining for 1 hour in 0.5% crystal violet in 20% ethanol.

## Stem cell culture

Human Y6 and Y1 iPS lines were obtained from Yale Stem Cell Center and New York Cell Stem Foundation respectively. Cells were verified as being pluripotent, having normal karyotype, mycoplasma free and cultured in feeder-free conditions on matrigel-coated plates with mTeSR Plus culture media (Stem Cell Technologies) and passaged using ReLeSR (Stem Cell Technologies).

## Teratoma formation

1×10^6^ Y6 iPSC cells were collected by collagenase treatment, and resuspended in 100 ml of DMEM/F12, collagen, and matrigel mix (2:1:1 ratio). Cells were intramuscularly injected into immunodeficient Rag2-/-GammaC-/- mouse. After 8 weeks, teratomas were harvested, fixed, and subjected to paraffin-embedding and haematoxylin and eosin (H/E) staining.

## G-band staining for Karyotype analysis

Y6 iPSC small clumps were seeded on glass slide pre-coated with Matrigel and fed with mTESR medium for three days. Then, medium were switched to DMEM basal medium supplemented with 10% of FBS for another 3 days. Then the slide was transferred to Yale Cytogenetic laboratory for G-band staining.

## Neural Progenitor Cell culture

Y6 iPS lines were differentiated to NPCs on matrigel-coated plates using the monolayer protocol of the StemDiff SMADI Neural Induction kit (Catalog # 08581, Stem Cell Technologies) for two passages and then maintained in StemDiff Neural Progenitor media (Catalog #05833, Stem Cell Technologies). Twelve-day NPCs were used for all experiments.

## Cerebral organoid culture

For preparation of embryoid bodies, 9000 single cells were seeded in each well of low attachment 96-well U-bottom plate and 10 μM Y-27632 ROCK inhibitor for one day. Cerebral organoids were generated following exactly the previously established protocol(Lancaster and Knoblich, 2014; Lancaster et al., 2013), using an orbital shaker for agitation.

## Immunostaining

Brains of infected mice and organoids were collected and fixed in 4% PFA. Samples were then dehydrated in a 30% sucrose solution. Cryostat sections were blocked in 0.1 M Tris-HCl buffer with 0.3% Triton and 1% FBS before staining. Slides were stained for IBA-1 (nb100-1028, Novus Biologicals), GFAP (ab4674, abcam and rabbit anti-SARS-CoV-2 nucleocapsid (GeneTex) Slides were mounted with Prolong Gold Antifade reagent (Thermo fisher). All slides were analyzed by fluorescence microscopy (BX51; Olympus).

Organoids were fixed in 4% PFA in a BSL3 facility then moved to 30% sucrose solution at 4°C for at least 24h. Organoids were then embedded in Tissue Freezing Medium (TFM) and cut into 20 μm sections using a cryostat and mounted on slides. After blocking and permeabilization with 0.25% Triton-X 100 and 4% donkey serum, sections were incubated overnight with primary antibody: ms anti-Pax6 (1:300, BD Pharmingen, #561462), rb anti-ACE2 (1:500, Abcam, ab15348), ms anti-Sox2 (1:500, Santa Cruz, sc-365823), rat anti-CTIP2 (1:500, Abcam, ab18465), rb anti-TBR1 (1:500, Abcam, ab31940), ms anti-MAP2 (1:500, Millipore, MAB3418), rb anti-SARS-CoV-2 nucleocapsid (1:250, GeneTex, GTX635679), rb anti-HIF1 alpha antibody (Genetex, GTX127309). TUNEL assay was performed using the Click-It Plus TUNEL Assay for In Situ Apoptosis Detection (Thermofisher, C10617) per the manufacturer’s instructions. Alexa Fluor 488, 555, 647 antibodies were applied for 1hr at room temperature (1:500) after three 10min PBS washes. To mark nuclei, DAPI (1:3000) was added to the secondary antibody incubation. Slides were then washed three times in PBS and then mounted with VectaShield Anti-Fade Mounting Medium. Images were acquired using a Zeiss LSM 880 confocal microscope (Carl Zeiss) and prepared using Fiji (NIH).

Neural progenitor cells on coverslips were fixed in 4% PFA for 15 minutes at room temperature in BSL3 conditions and washed 3x with PBS. After blocking and permeabilization with 0.25% Triton-X 100 and 10% donkey serum, coverslips were incubated overnight with primary antibody as described above. TUNEL assay was performed as described above. Alexa Flour 488, 555, and 647 antibodies were applied for 1hr at room temperature (1:500) after three 10min PBS washes. Coverslips were then washed three times in PBS and then mounted on slides with Prolong Diamond Anti-Fade Mountant with DAPI (Thermofisher). Images were acquired using a Zeiss LSM 880 confocal microscope (Carl Zeiss) and prepared using Fiji (NIH).

Human Formalin-fixed Paraffin sections were heated for 30 minutes at 60°C and treated with xylenes followed by rehydration in decreasing concentrations of ethanol (100%, 90%, 80% 70%). Antigen retrieval was performed using a pressure cooker (Biocare Decloaking Chamber) by boiling in sodium citrate (pH 6.0) for 20 minutes at 115 degrees. After blocking and permeabilization with 0.25% Triton X 100 and 4% donkey serum, sections were incubated overnight with primary antibody: rb anti-ACE2 (1:500, ab15348, Abcam), ms anti-NeuN (1:500, ab104224, Abcam). Alexa Fluor 488, 555 antibodies were applied for 1 hr at room temperature (1:500) after three 10min PBS washes. To mark nuclei, DAPI (1:3000) was added to the secondary antibody incubation. Slides were then washed three times in PBS to remove detergent and treated with TrueBlack Lipofuscin Autofluorescence Quencher (Biotium) according to the manufacturer’s instructions. Sections were mounted with VectaShield mounting medium. Images were acquired using a Zeiss LSM 880 confocal microscope (Carl Zeiss) and prepared using Fiji (NIH).

## Electron microscopy

Organoids were fixed using 2.5% glutaraldehyde in 0.1M phosphate buffer, osmicated in 1% osmium tetroxide, and dehydrated in ethanol. During dehydration, 1% uranyl acetate was added to the 70% ethanol to enhance ultrastructural membrane contrast. After dehydration the organoids were embedded in Durcupan and 70 nm sections were cut on a Leica ultramicrotome, collected on Formvar coated single-slot grids, and imaged on a Tecnai 12 Biotwin electron microscope (FEI).

## Single-cell RNA-seq

Organoids were collected and single cell suspensions were made using a papain dissociation system (Worthington Biochemical Corporation). Single-cell suspensions were loaded onto the Chromium Controller (10x Genomics) for droplet formation. scRNA-seq libraries were prepared using the Chromium Single Cell 3′ Reagent Kit (10x Genomics. Samples were sequenced on the on the NovaSeq.

R v.3.4.2 (R Core Team 2013) was used for all statistical analysis. Sequencing results were demultiplexed into Fastq files using the Cell Ranger (10x Genomics, 3.0.2) mkfastq function. Samples were aligned to GRCh38 10x genome. The count matrix was generated using the count function with default settings. Matrices were loaded into Seurat v.3. 1.5 for downstream analysis. Cells with less than 200 unique molecular identifiers, or high mitochondrial content were discarded. Using FindIntegrationAnchors and IntegrateData functions, all libraries were integrated into a single matrix. Principal component values that were statistically significant were identified, and a cut-off point was determined using the inflection point after using the PCElbowPlot function. Clusters were determined using the RunUMAP, FindNeighbors, and FindClusters functions on Seurat. Cells were considered infected if transcripts aligned to viral ORF1ab, Surface glycoprotein (S), ORF3a, Envelope protein (E), Membrane glycoprotein (M), ORF6, ORF7a, ORF8, Nucleocapsid phosphoprotein (N) or ORF10. Differentially expressed genes from the FindMarkers function were used to perform PANTHER-GO statistical over-representation tests for up-regulated and downregulated genes in each condition shown in Figure 2F and G. Gene lists for Figure 2H were obtained from GSEA databases (gsea-msigdb.org/gsea/index.jsp).

## Immunohistochemistry

Paraffin sections were heated for 30 minutes at 60°C and treated with xylenes followed by rehydration in decreasing concentrations of ethanol (100%, 90%, 80%, 70%). Antigen retrieval was performed by boiling in sodium citrate (pH 6.0) for 15 minutes and peroxidase activity was blocked with hydrogen peroxide for 10 minutes. Blocking was performed in 2.5% normal horse serum (Vector Laboratories) and incubated in primary antibody overnight at 4°C. Mouse anti-SARS-CoV-2 spike antibody (clone 1A9, GeneTex GTX632604) was used at a dilution of 1:400. Secondary antibody and detection reagents from the VECTASTAIN Elite ABC-HRP Kit (Vector Laboratories PK-7200) were used according to manufacturer instructions. Sections were counterstained with Hematoxylin QS (Vector Laboratories H-3404), dehydrated in increasing concentrations of ethanol, cleared with xylenes, and mounted with VectaMount permanent mounting medium (Vector Laboratories H-5000).

## ACE2 blocking

For ACE2 blocking assays, Organoids were preincubated with anti-ACE2 Antibodies (AF933, R&D Systems) or isotype antibodies at a concentration of 100ug/mL for 1 hour at 4 degrees. Organoids were then infected with SARS-CoV-2 as described above.

## CSF neutralization assay

Excess CSF was obtained from a hospitalized patient with COVID-19 who underwent clinical LP, and from a healthy control volunteer. CSF was centrifuged to isolate cell-free supernatant, which was used for ELISA and neutralization assays as follows:

### Enzyme-linked immunosorbent assay

ELISAs were performed as previously reported (Israelow et al., 2020). In short, Triton X-100 and RNase A were added to serum samples at final concentrations of 0.5% and 0.5mg/ml respectively and incubated at room temperature (RT) for 3 hours before use to reduce risk from any potential virus in serum. 96-well MaxiSorp plates (Thermo Scientific #442404) were coated with 50 μl/well of recombinant SARS Cov-2 S1 protein (ACROBiosystems #S1N-C52H3-100ug) at a concentration of 2 μg/ml in PBS and were incubated overnight at 4 °C. The coating buffer was removed, and plates were incubated for 1h at RT with 200μl of blocking solution (PBS with 0.1% Tween-20, 3% milk powder). Serum was diluted 1:50 in dilution solution (PBS with 0.1% Tween-20, 1% milk powder) and 100μl of diluted serum was added for two hours at RT. Plates were washed three times with PBS-T (PBS with 0.1% Tween-20) and 50μl of mouse IgG-specific secondary antibody (BioLegend #405306, 1:10,000) diluted in dilution solution added to each well. After 1h of incubation at RT, plates were washed three times with PBS-T. Samples were developed with 100μl of TMB Substrate Reagent Set (BD Biosciences #555214) and the reaction was stopped after 15 min by the addition of 2 N sulfuric acid.

### Neutralization assay

Virus for infection was preincubated with 500μL of healthy or COVID19 CSF in 37 degrees for one hour before infection of organoids. The Organoid culture was supplemented with an additional 500μL of CSF after infection until the organoid was collected for imaging.

## Human subjects

Research participants were enrolled at Yale University through Human Investigation Committee Protocols HIC#2000027690 and HIC #1502015318. The Institutional Review Board at Yale approved the protocols, and informed consent was obtained from all participants.

Post-mortem COVID-19 brain tissues were obtained from COVITIS Biobank (Assistance Publique Hopitaux de Paris, Paris, France).

## Statistical analysis

No statistical methods were used to predetermine sample size. The investigators were not blinded during experiments and outcome assessment, but outcome assessment was additionally evaluated by animal technicians and vets blinded to the study. Survival curves were analyzed using a log-rank (Mantle-Cox) test. For other data, normally distributed continuous variable comparisons used a two-tailed unpaired Student’s *t*-test or paired Student’s *t*-test with Prism software.

## Acknowledgements

We thank Yale Environmental Health and Safety (EHS) department for allowing safe working environments with the SARS-CoV-2 virus. We also thank the patient donors, and clinicians who helped with the collection of CSF for neutralization assays. We also thank the members of YCGA who have helped with all aspects of sequencing. Finally, we thank the members of the Iwasaki lab for insightful discussions regarding the project. This study was supported by awards from National Institute of Health grants, R01AI157488 (AI, SFF), R01NS111242 (AI) T32GM007205 (MSTP training grant), F30CA239444 (ES), 2T32AI007517 (BI), K23MH118999 (SFF) Women’s Health Research at Yale Pilot Project Program (AI, AR), Fast Grant from Emergent Ventures at the Mercatus Center (AI, ES, CBW), Mathers Foundation (AR, CBW, AI), and the Ludwig Family Foundation (AI, AR, CBW). A.I. is an investigator of the Howard Hughes Medical Institute.

## Author Contributions

E. S, C.Z., K.B., and A.I. planned the project and analyzed data. E.S., C.Z. and A.I. wrote the manuscript. E.S., C.Z., B.I., A.L., A.V.P., S.S., performed experiments. P.L., O.E.W., F.L., Y.D., K.S.B., Y.Y., G.W., J.H., E.N., J.W., M.M.A, assisted with experiments. E.L., S.A.J.K., K.Z., A.R., C.B.W., T.L.H., I.P., N.R., S.H., J.L.T., A.H., D.S., A.L., S.F.F., and K.B. provided expertise and materials for analysis of data. A.V.P., S.S., N.R., performed iDISCO+ imaging and analysis of mice. E.L., S.A.J.K., K.Z., I.P., S.H., J.L.T., A.H., D.S., provided human samples and analysis of images. A.I. and K.B. supervised the project and secured funding.

## Declaration of Interests

None of the authors declare interests related to the manuscript.

## Materials & Correspondence

Correspondence and material requests should be addressed to

Requests for scripts, code and raw data used in this study should be addressed to

---

**Figure S1:**
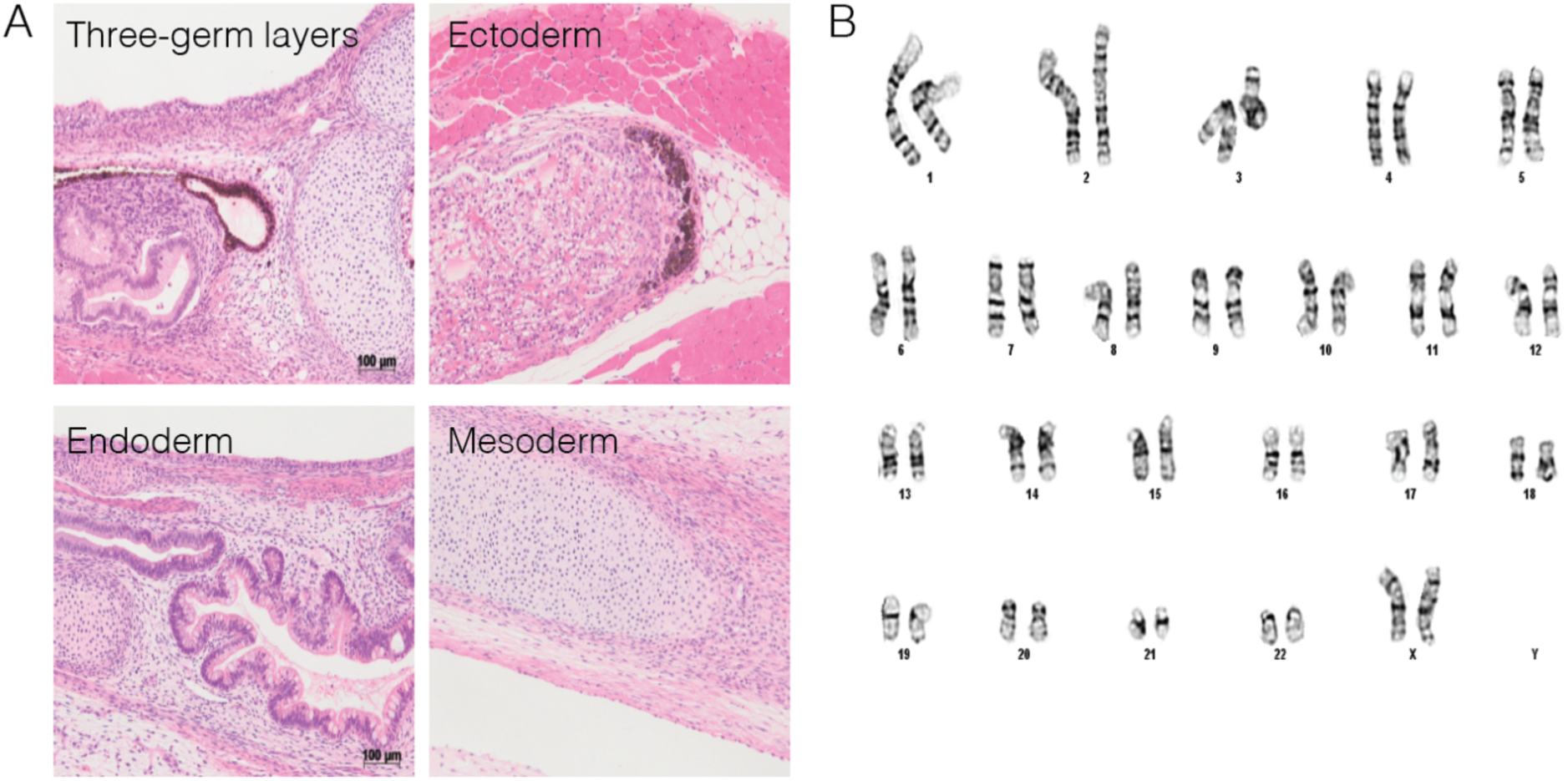
Pluripotency of Y6 line. (**A**) Sample images of cell types of three germ-layers in teratomas following transplantation to Rag2-/-GammaC-/- mice. Scale bar 100um. (**B**) Sample image showing normal karyotype for Y6 line.

**Figure S2:**
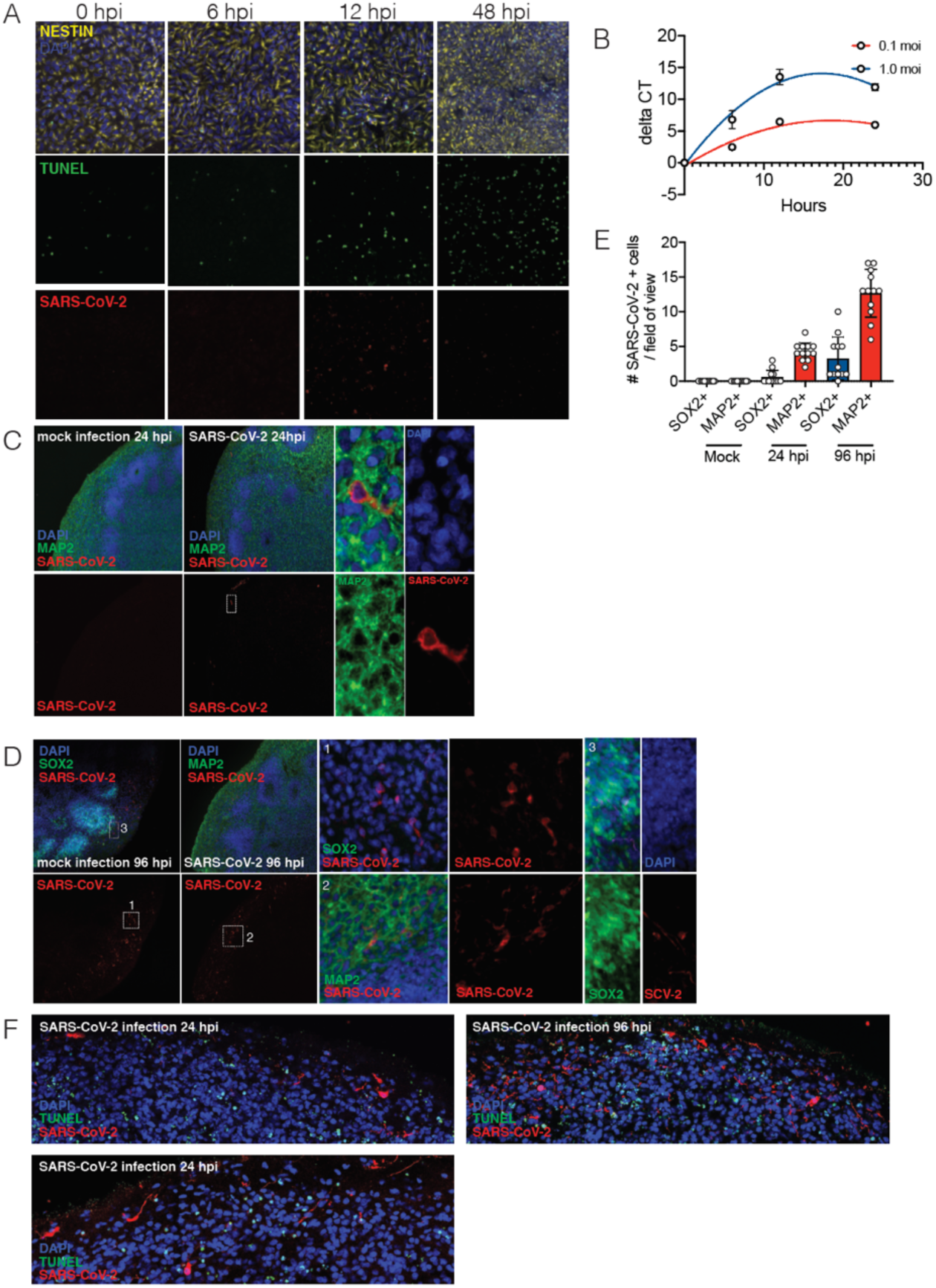
SARS-CoV-2 infection in hPSC-derived neural progenitor cells and 9-week organoids. (**A**) Representative images of immunostaining of hNPCs with TUNEL staining, anti-Nestin, and anti-SARS-CoV-2 nucleocapsid antibody at 0, 6, 12, and 48 hours after infection. (**B**) RT-PCR amplification of COVID-genome from infected hNPC cells. (**C**) Sample images of immunostaining of 9-week organoids with DAPI, anti-MAP2, and anti SARS-CoV-2 ab at 24 hpi after SARS-CoV-2 or mock infection. Note the perinuclear and neuritic staining of SARS-CoV-2 in the MAP2+ cell. Dashed white box corresponds to SARS-CoV-2+ and MAP2+ single cell in 24 hpi organoid. (**D**) Sample images of immunostaining of 9-week organoids with DAPI, anti-MAP2 or anti-Sox2, and anti-SARS-CoV-2 ab at 96 hpi after SARS-CoV-2 or mock infection. Dashed white box “1” corresponds to “1 panel” showing magnified SARS-COV-2+/SOX2-cell in 96 hpi organoid. Dashed white box “2” corresponds to “2 panel” showing SARS-COV-2+ and MAP2+ cell in 96 hpi organoid. “3” panel shows SOX2+/SARS-CoV-2+ cell in 96 hpi organoid. (**E**) Quantification of number of SARS-CoV-2 + cells / field of view that are either SOX2+ or MAP2+ in mock, 24 hpi, and 96 hpi, organoids. (**F**) Organoids were stained with TUNEL to evaluate cell death at 24hpi and 96hpi. N=4 organoids per condition, 12 cortical regions from two iPSC lines.

**Figure S3:**
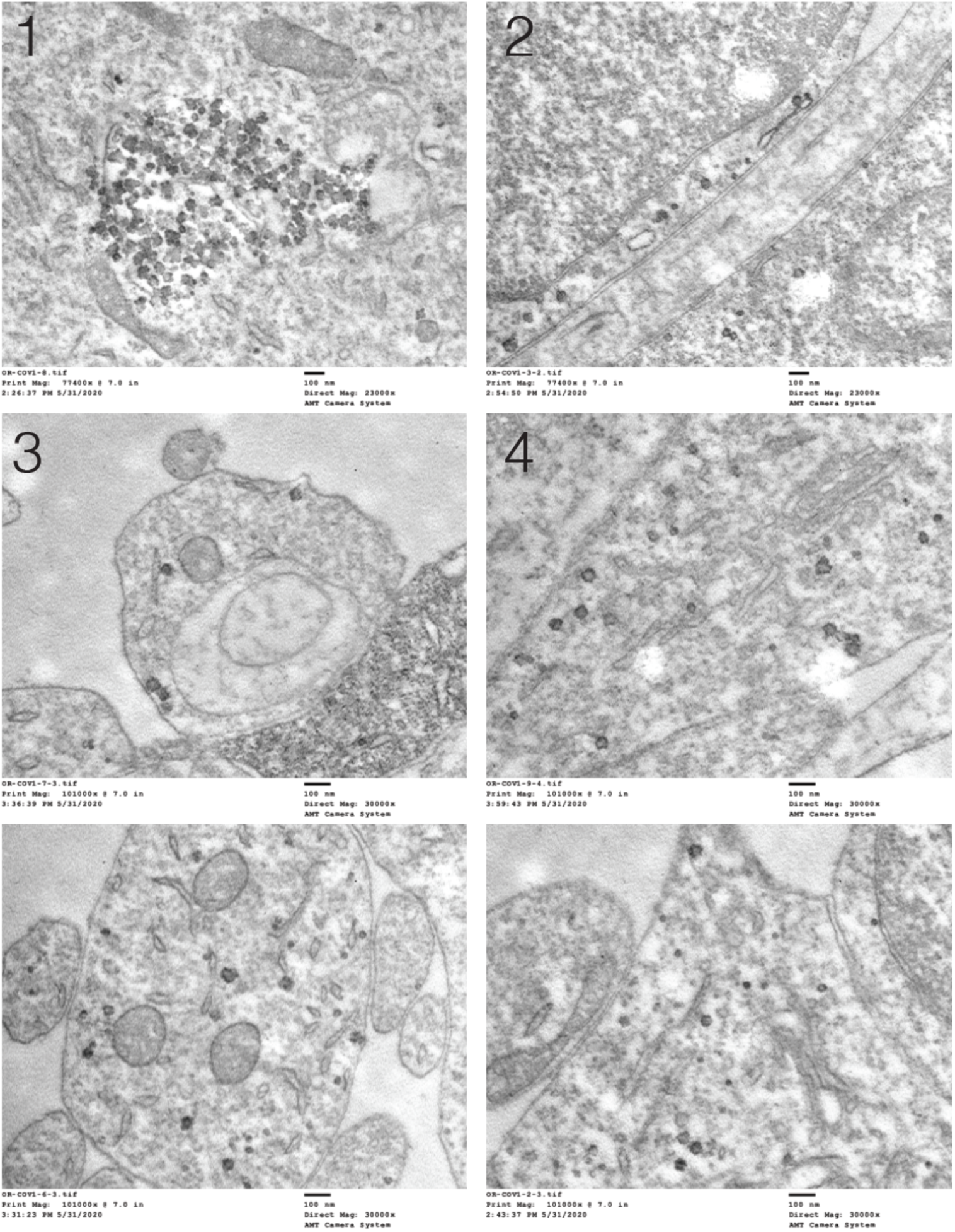
Electron Microscopy of SARS-CoV-2 infection. Uncropped images of electron microscopy of infected brain organoids from Fig. 1D.

**Figure S4:**
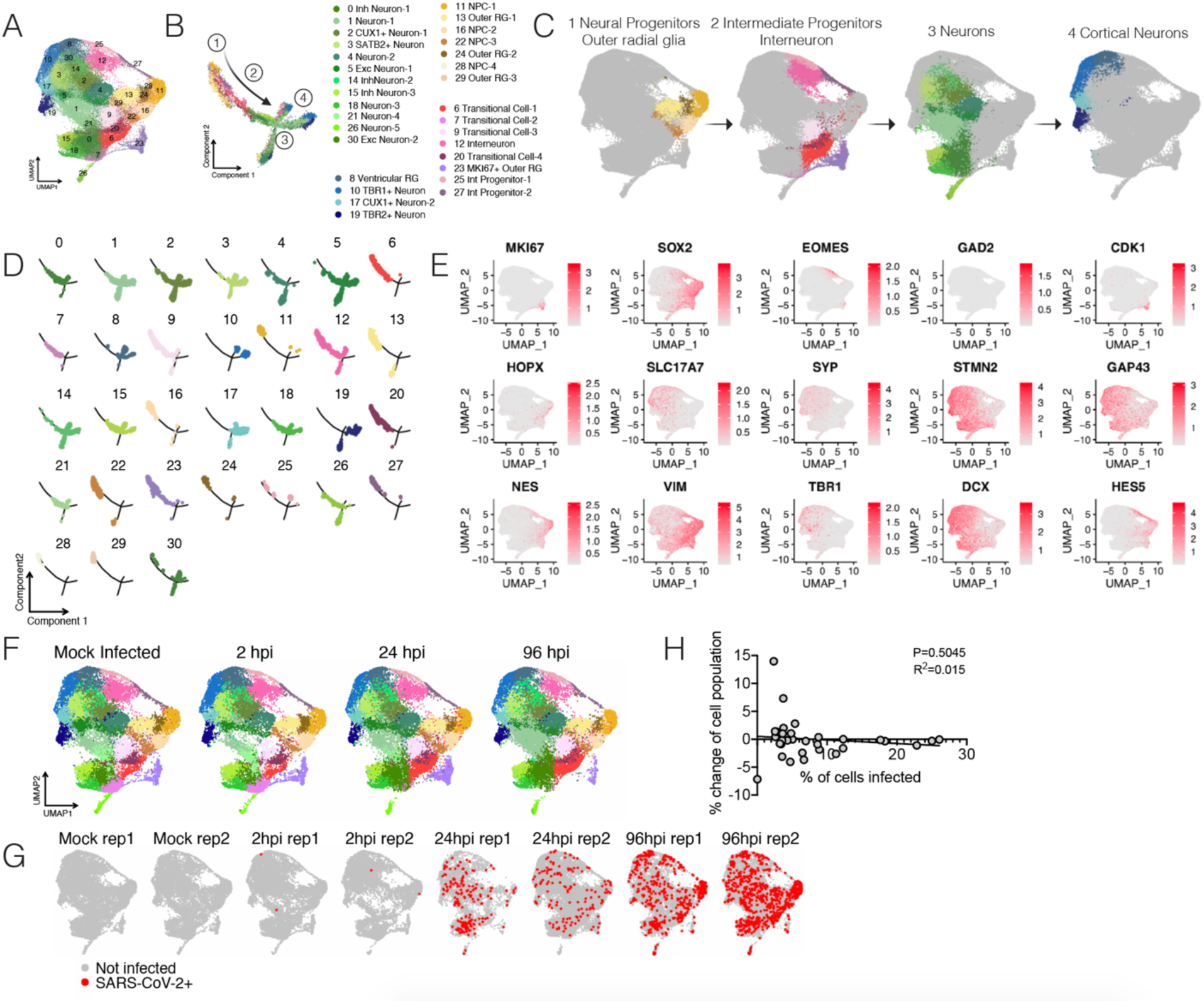
Single cell RNA-seq of SARS-CoV-2 infected organoids. (**A**) UMAP projection of cells from single cell sequencing. (**B**) Monocle trajectory analysis resulted in four distinct states of cells from the organoid. (**C**) The four major clusters consisted of 1. Neural progenitor, outer radial glia like cells, 2. Intermediate progenitor, interneurons, 3. Neurons, and 4. Cortical neurons. (**D**) Monocle projection of individual clusters. (**E**) Heatmap of commonly used genes for identification of cell subtypes in human brain organoids. (**F**) UMAP projection of organoids depending on infection status. (**G**) UMAP heatmap of SARS-CoV-2 transcript + cells separated by infection status. (**H**) Correlation between % change of cell population versus % of cells infected in each cluster.

**Figure S5:**
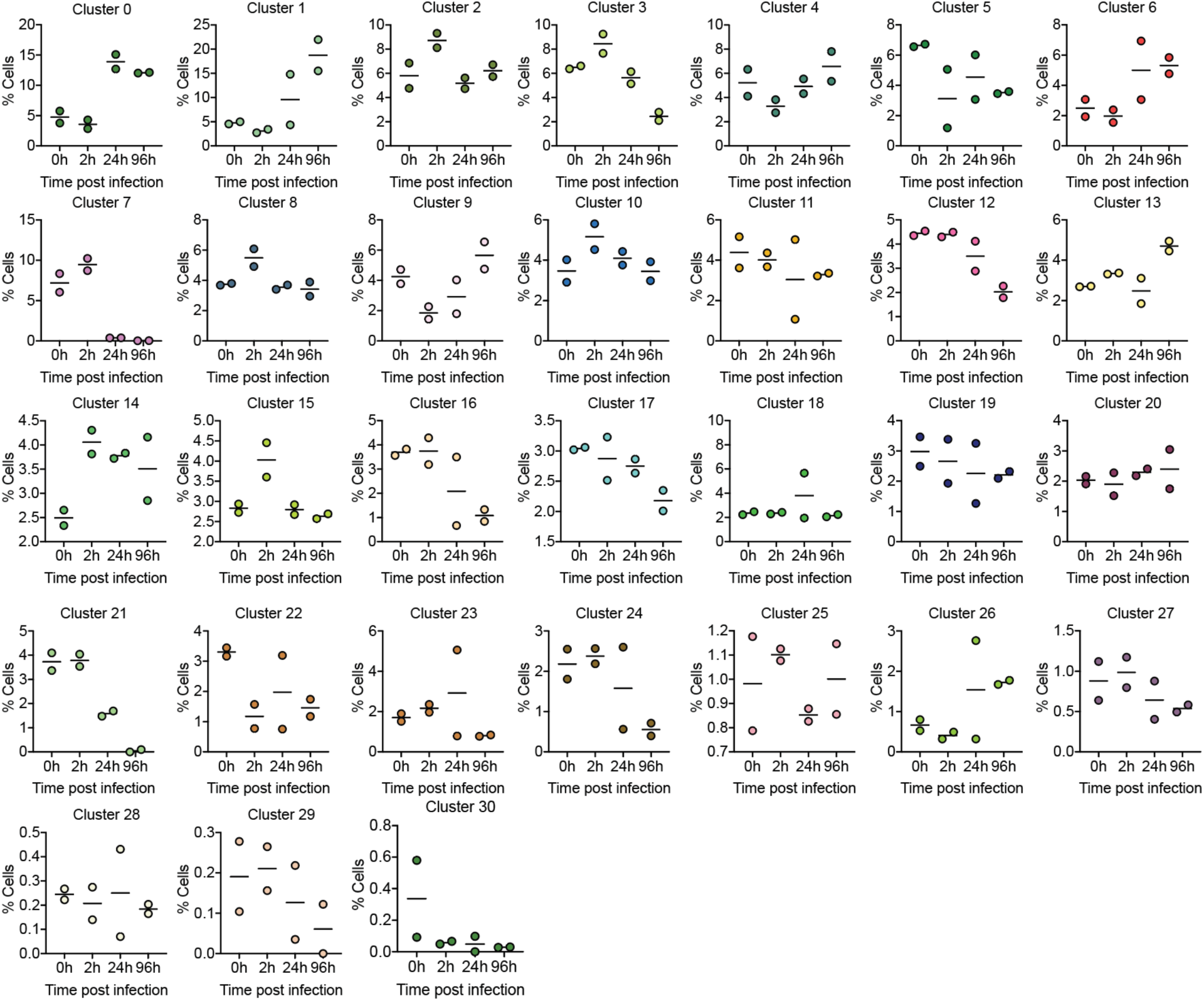
Frequency of each cell cluster from single cell RNA-seq. Graphs display the % frequency of each cluster during a given infection status, 0, 2, 24 and 96 hours post infection.

**Figure S6:**
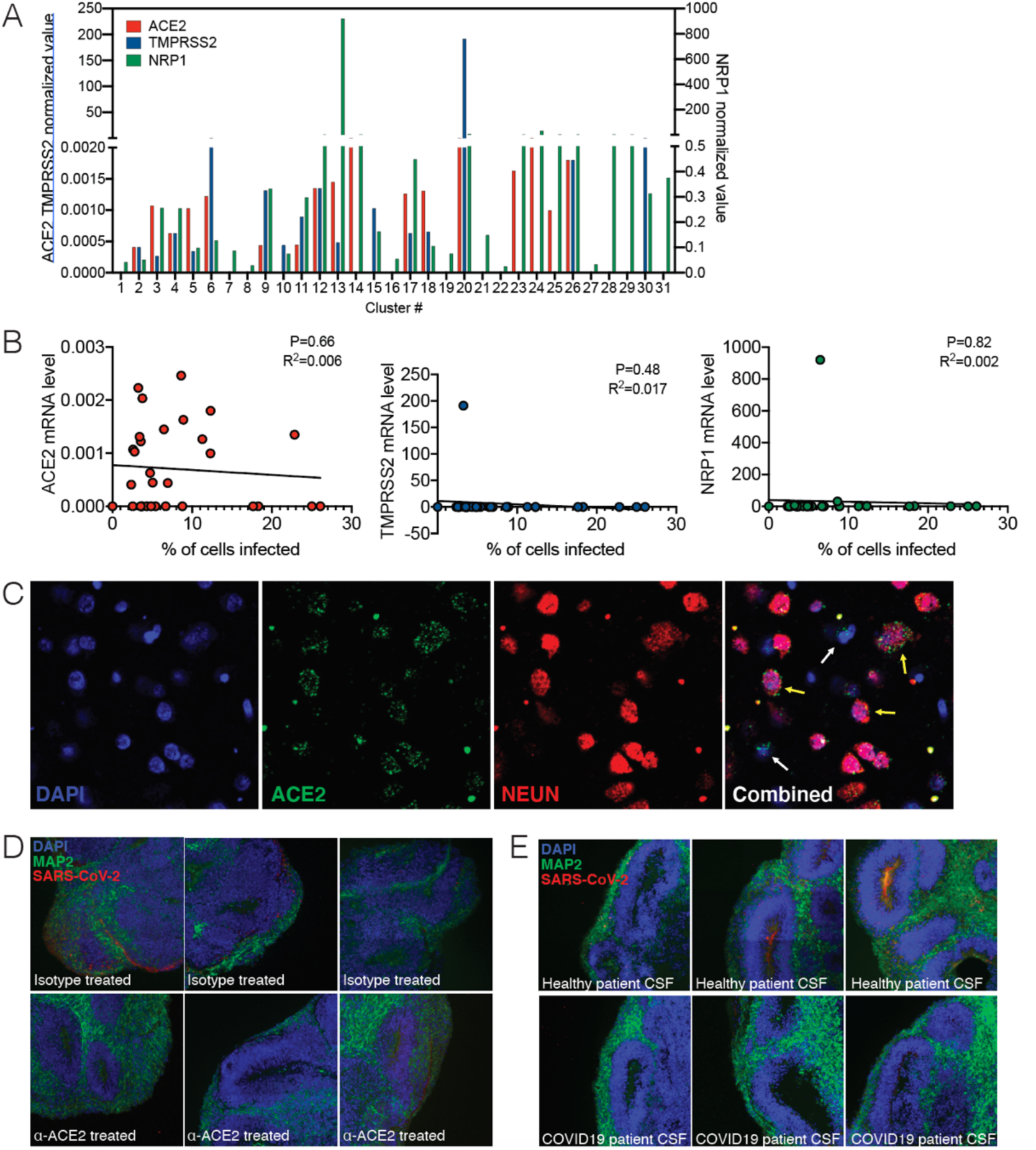
SARS-CoV-2 infection depends on hACE2. (**A**) Expression levels of ACE2, TMPRSS2, and NRP1 from single cell RNA-seq data. (**B**) Correlation of ACE2, TMPRSS2 and NRP1 expression to % of cells infected in each cluster. (**C**) Imaging of post-mortem COVID-19 patient brains stained with ACE2 and NEUN. (**D**) Uncropped images of ACE2-antibody treated organoids shown in 3B. (**E**) Uncropped images of CSF-treated organoids shown in 3E. All experiments were performed with unique organoid n of 4 per condition, with images from n = 12 cortical regions with two IPSC lines.

**Figure S7:**
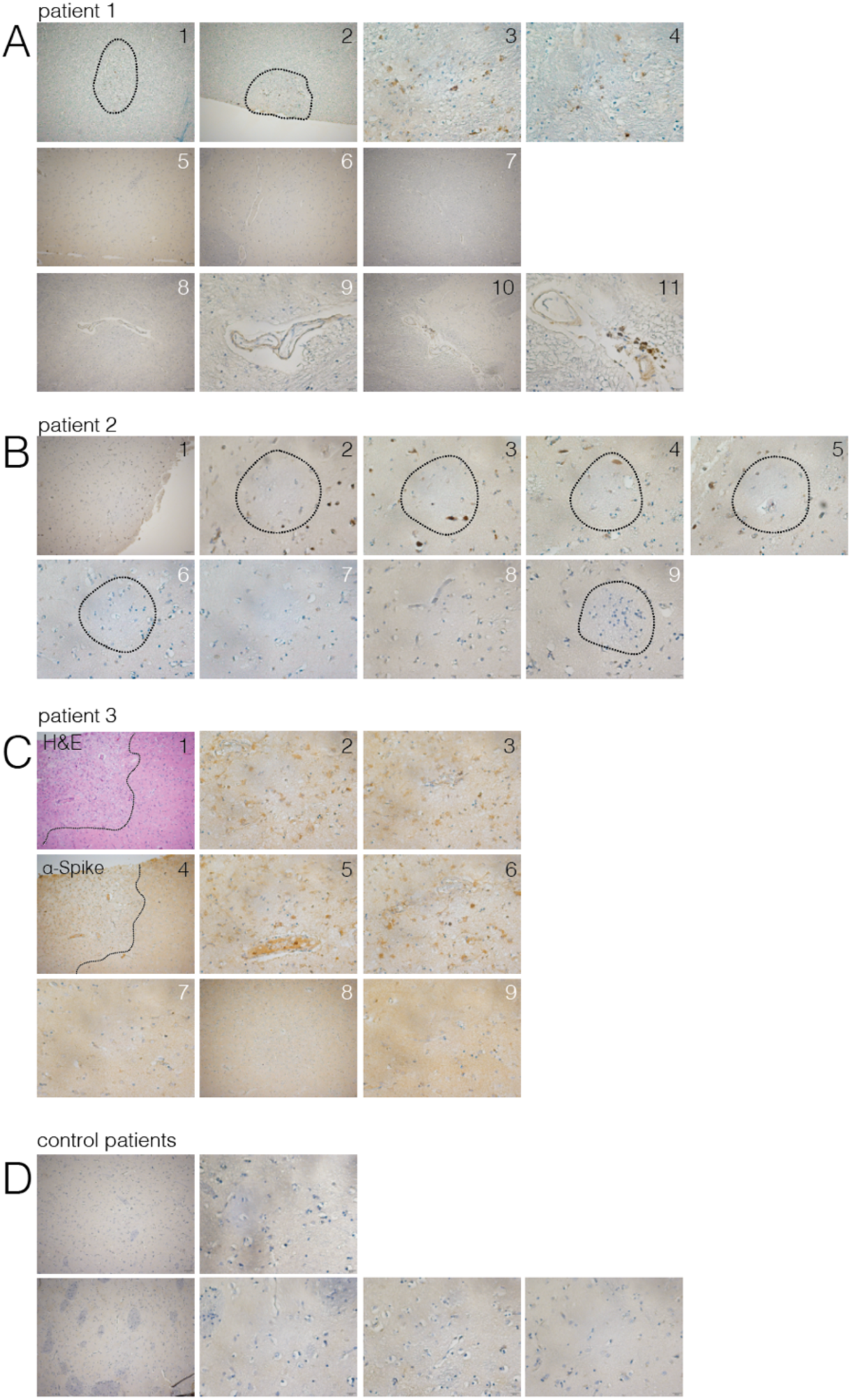
Evidence of neuroinvasion in post-mortem COVID-19 patient brains (Caudate). FFPE sections of brain tissue from COVID-19 patients were stained using H&E or anti-SARS-CoV-2-spike antibody. (**A**) Images of regions of the caudate of patient 1. White numbers indicate unaffected regions, and black numbers indicate regions with infected cells. Dotted lines around 1 and 2 indicate ischemic infarcts with virus staining. (**B**) Images of regions of the caudate of patient 2. White numbers indicate unaffected regions, and black numbers indicate regions with infected cells. Dotted circles indicate ischemic infarcts. (**C**) Images of regions of the caudate of patient 3. White numbers indicate unaffected regions, and black numbers indicate regions with infected cells. Dotted lines around 1 and 4 indicate ischemic infarcts with virus staining. Images 1, 4 and 5 are shown in main figure 4. (**D**) Example images from control patient brains.

**Figure S8:**
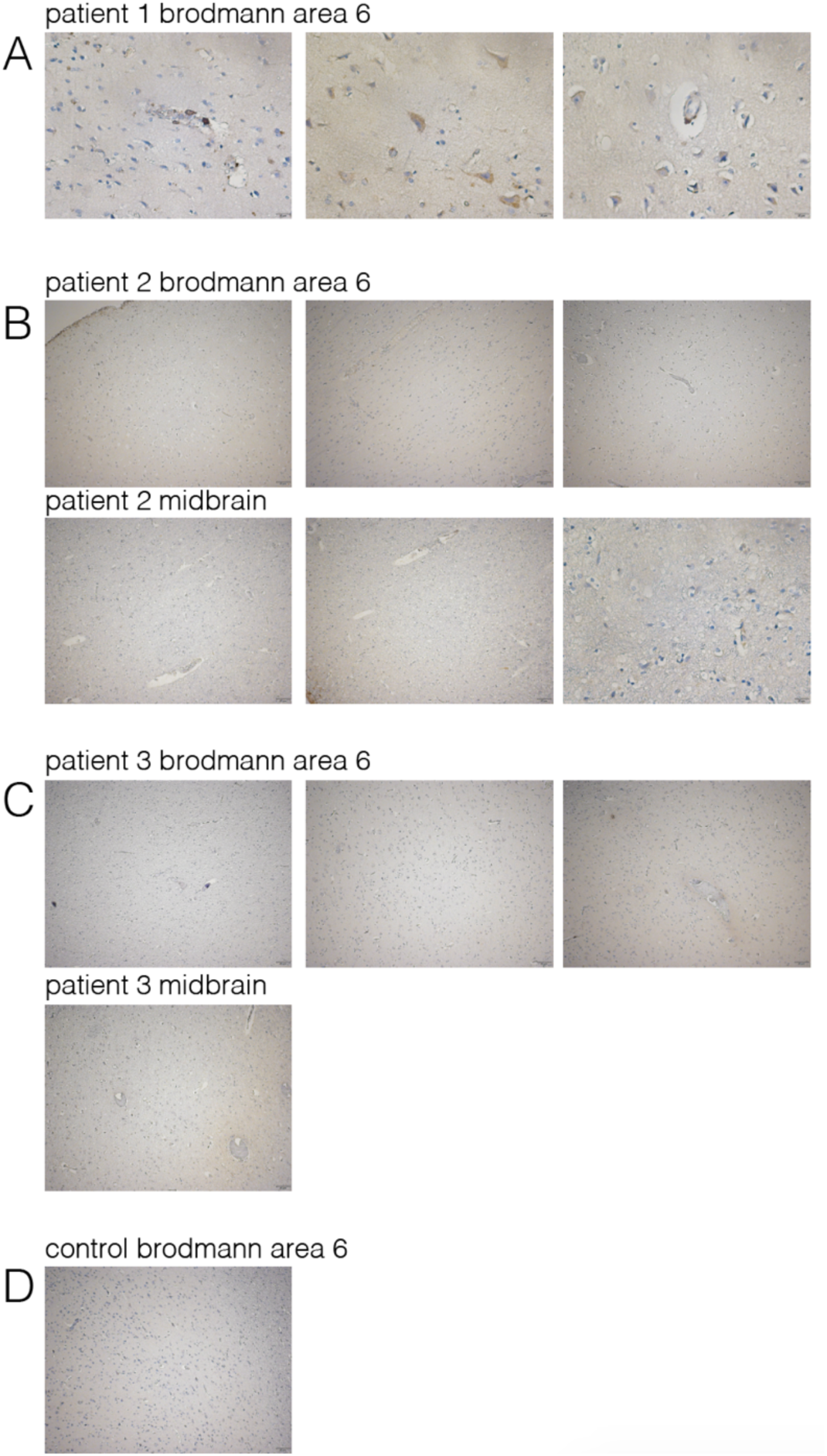
Evidence of neuroinvasion in post-mortem COVID-19 patient brains (Brodmann area 6 and midbrain). FFPE sections of brain tissue from COVID-19 patients were stained using anti-SARS-CoV-2-spike antibody. (**A**) Images of regions of Brodmann area 6 of patient 1. Images are also shown in main figure 4. (**B**) Images of regions of patient 2. (**C**) Images of regions of the caudate of patient 3. (**D**) Example image from control patient brains.

**Figure S9:**
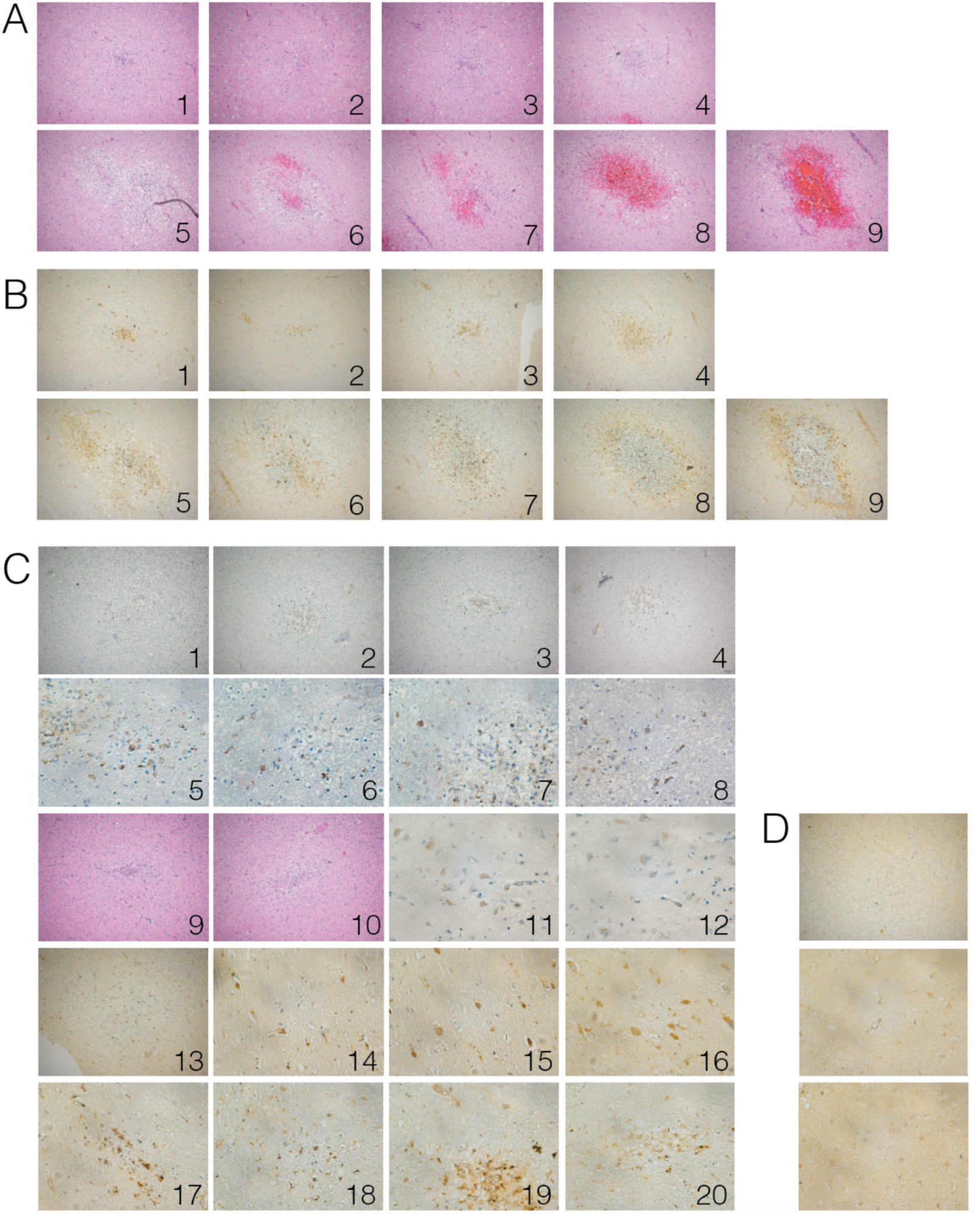
Evidence of SARS-CoV-2 neuroinvasion associated ischemic infarcts. FFPE sections of brain tissue from COVID-19 patients were stained using H&E or anti-SARS-CoV-2-spike antibody. (**A**) H&E images of ischemic infarcts at different stages (1 being earliest to 9 being latest). (**B**) SARS-CoV-2 stained images of ischemic infarcts at different stages (1 being earliest to 9 being latest). Each number corresponds to H&E image found in **A**. (**C**) Images of SARS-CoV-2 positive regions in brains of COVID-19 patients. (**D**) Example image from control patient brains.

**Figure S10:**
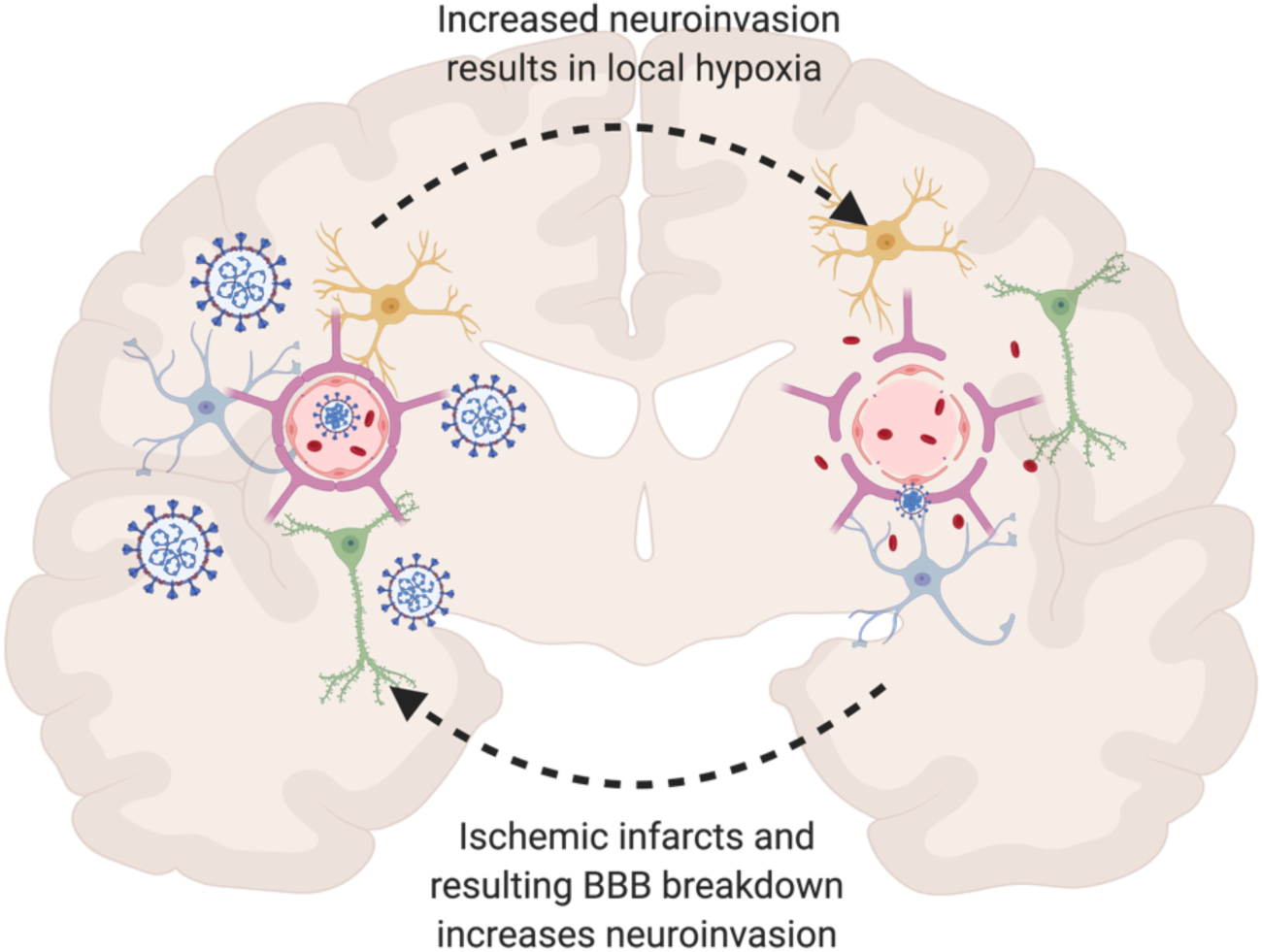
Schematic of hypothesized consequences of SARS-CoV-2 neuroinvasion.

**Supplementary Movie 1:**
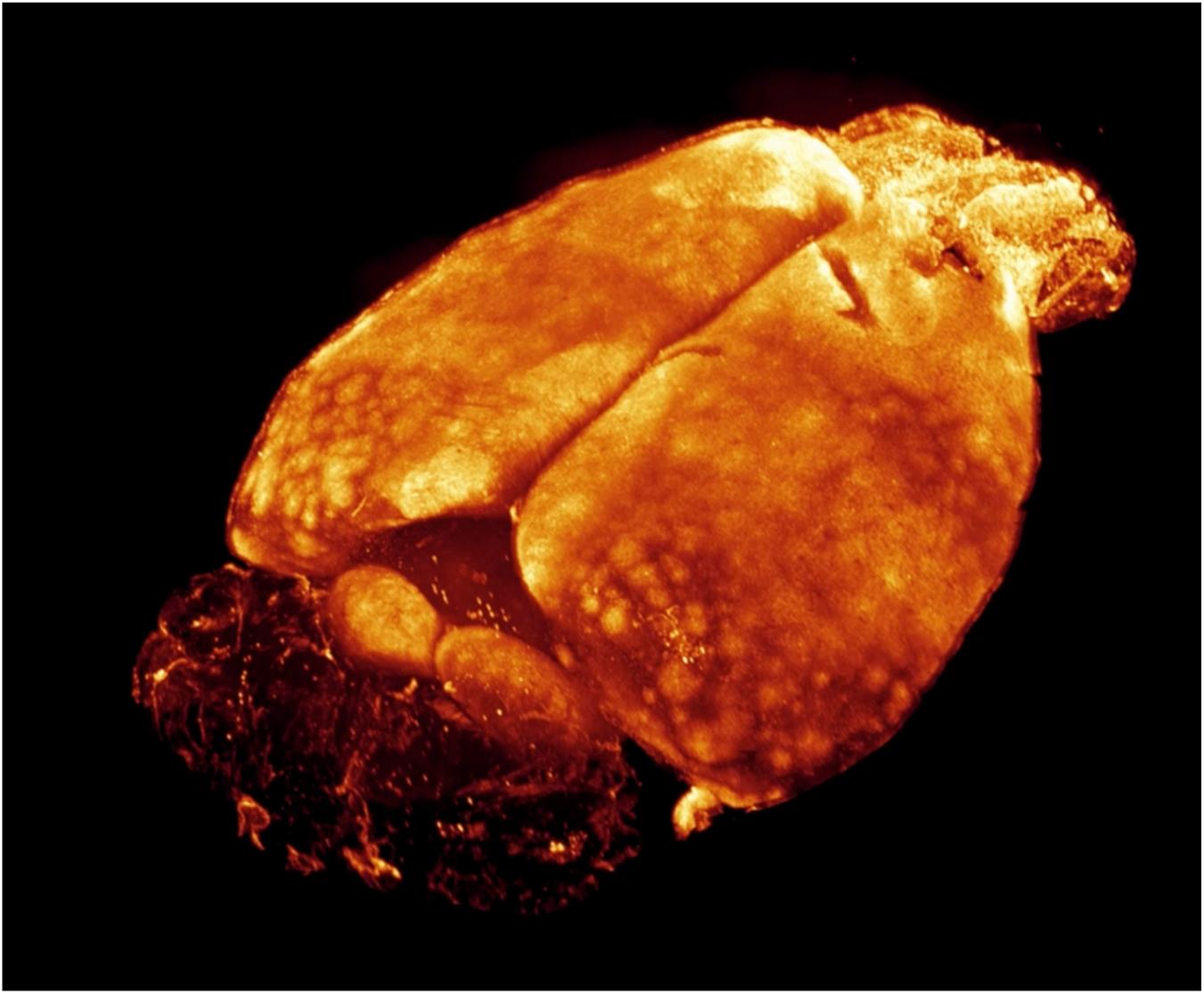
Whole brain view of SARS-CoV-2 infection. iDISCO+ whole brain immunolabeling against the nucleocapsid protein of SARS-CoV2 7 days after an intranasal infection

**Supplementary Table 1:**
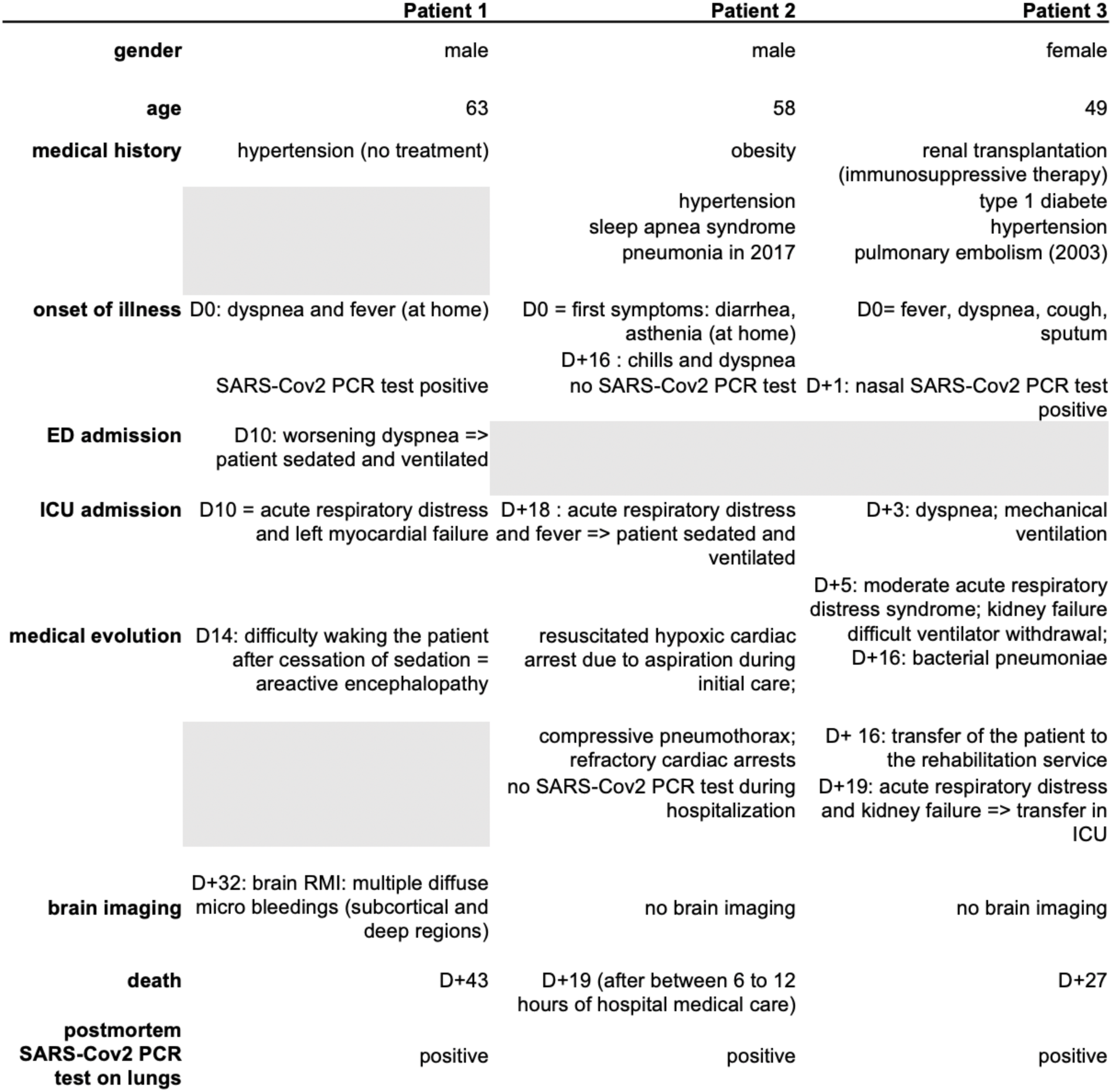
Clinical characteristics of COVID-19 patients.

See attached excel file

**Supplementary Table 2: Normalized gene count of clusters found in SARS-CoV-2**

## Notes

### Competing Interest Statement

The authors have declared no competing interest.

### Summary of Updates

This version contains brain autopsy SARS-CoV-2 infection data.

